# Phylogenetically distant but cohabiting: Fungal communities of fine roots in *Diphasiastrum complanatum*, *Pinus sylvestris*, and *Vaccinium myrtillus* in a Lithuanian pine forest

**DOI:** 10.1101/2025.01.12.631741

**Authors:** Kristina Kuprina, Moana Wirth, Maria Sanchez Luque, Heike Heklau, Radvilė Rimgailė-Voicik, Manuela Bog, Martin Schnittler

## Abstract

Throughout evolution, distinct plant lineages independently established mutualistic relationships with various fungal taxa. However, the extent to which these relationships are conserved across different plant and fungal lineages remains unclear. In this study, we compared fungal communities associated with the fine roots of three phylogenetically distant yet cohabiting plant species: *Diphasiastrum complanatum*, a member of lycophytes, the most basal extant vascular plant lineages; *Pinus sylvestris,* a gymnosperm; and *Vaccinium myrtillus*, an angiosperm, an evolutionary relatively young lineage. To minimize environmental variability, fine roots of three species were collected from each of 19 five-square-meter plots within a Scots pine forest in Lithuania. Using metabarcoding and microscopic techniques, we observed significant differences in the fungal community composition and diversity among the three plant species. We detected no signs of arbuscular mycorrhiza in any species. Samples of *D. complanatum* showed significantly higher taxonomical diversity, while *P. sylvestris* showed lowest diversity, with ectomycorrhizal fungi being most abundant. Samples of *V. myrtillus* had a prevalence of putative ericoid mycorrhiza taxa, classes Sebacinales and Trechisporales, likely forming hyphal coils detected through microscopy. In contrast, no mycorrhiza was detected in *D. complanatum* sporophytes. This, along with the presence of well-developed root hairs and similarity to the fungal community inhabiting soil suggests a low dependency on mycorrhizal associations and a more opportunistic fungi-plant relationship. This is the first study of fungi associated with the sporophytes of *D. complanatum.* Our findings provide valuable insights into the complex interactions between fungi and plants from diverse phylogenetic lineages in natural environments.

## 1. Introduction

The widespread symbiotic relationships between fungi and plants profoundly influence ecosystem structure and function (Grime et al. 1987; Clay and Schardl 2002; Stone et al. 2004; Frank and Trappe 2005; Smith and Read 2008; Brundrett and Tedersoo 2018). Endophytic and mycorrhizal fungi are key examples of these relationships, they coexisted with plants for over 400 million years and play an important role in driving their evolutionary trajectory (Remy et al. 1994; Krings et al. 2007; Redecker et al. 2000). Endophytic fungi inhabit plant tissues internally remaining asymptomatic and not causing visible disease (Petrini 1991). In exchange for nutrition and protection against abiotic and biotic stresses provided by the host, they can enhance the host’s fitness by increasing plant resistance to drought, heat, and salinity, and by producing secondary metabolites that protect against pathogens (Rodriguez et al. 2004; Rho et al. 2017). Mycorrhizal fungi, in turn, colonize not only plant roots but also the rhizosphere. This mutualistic association benefits both partners: the fungi supply the plant with organic or inorganic phosphates and nitrogen compounds, while the plant provides carbon for the fungi. As a result, approximately 80% of plant species form mycorrhizal relationships and thrive best with specific fungal partners (Wang and Qiu 2006; Park and Eom 2007; Smith and Read 2008; Delavaux et al. 2017).

There are several distinct types of mycorrhizal associations, each with a unique physiology, resource usage and specific fungal taxa (Wang and Qui 2006; Smith and Read 2008; Brundrett and Tedersoo 2018; Rimington 2020). Arbuscular mycorrhiza (AM) evolved first around 400 Mya through associations between early land plants and Glomeromycota and Endogonomycetes fungi, improving carbon and nutrient exchange via intracellular structures called arbuscules (Remy et al. 1994; Orchard et al. 2017; Strullu-Derrien et al. 2018). Ectomycorrhiza (ECM) evolved independently around 200 Mya from saprotrophic ancestors within various fungal lineages (Tedersoo and Brundrett 2017; Tedersoo and Smith 2013). The first detection of ectotrophic mycorrhiza with angiosperms dates back to a fossil from Tertiary amber (52 Mya; Beimforde et al. 2011). ECM fungi develop a hyphal mantle around the fine roots of a plant, where nutrient and carbon exchange take place within the Hartig net, a hyphal network that develops between the root’s epidermal and cortical cells. Due to their extensive hyphal systems, ECM fungi scavenge more effectively and at greater distances from the host roots compared with arbuscular mycorrhizal (AM) fungi (Teste et al. 2020). It was also shown that ECM symbiosis exhibits greater host specificity than AM symbiosis, which may be attributed to the enhanced dispersal capabilities of ECM fungi (Smith and Read 2008). Ericoid mycorrhizal (ERM) symbiosis originated around 117 Mya in Ericales initially with Ascomycota fungi, followed by Basidiomycota fungi (Schwery et al. 2015). At present, it is the least explored type of mycorrhiza (Vohník 2020). ERM fungi form intracellular hyphal coils in the plant roots and can also act as non-symptomatic endophytes in non-ericoid plants and as saprotrophs in soil and organic matter (Kohout 2017). Although ERM has been considered the most host-specific of all mycorrhizae, an increasing number of studies highlights its surprisingly promiscuous nature (Perotto et al. 2002; Kjøller et al. 2010; Walker et al. 2011).

Recent findings depict plant-fungus interactions as a complex and dense network, linking different plant taxa to fungi from various lineages (Simard et al. 2012; Verbruggen et al. 2012). The main fungal phylogenetic and functional groups are present in all ecosystems worldwide, though their relative proportions vary among biomes (Tedersoo et al. 2012). Plant-fungal associations are not random; plants select specific fungal symbionts, and fungi show preferences for certain host plants (Kennedy et al. 2003; Kiers et al. 2011; Bruns et al. 2002; Tedersoo et al. 2008; Öpik et al. 2009; Walker et al. 2011; Davison et al. 2011). At the same time, phylogenetically distant plant lineages can exhibit substantial overlap in fungal taxa (Öpik et al. 2009; Toju et al. 2013; Perez-Lamarque et al. 2022). These findings help to elucidate how these preferences influence plant and fungal species composition and contribute to the formation of different biomes.

Conducting studies of the root-associated fungi (RAF) of plants in natural biomes presents multiple challenges. One major issue is the close association between roots and soil, making it difficult to distinguish mutualistic fungi from free-living or transient fungi. At current, no method can reliably reveal whether adjacent growing plants are connected via a common mycorrhizal network within a natural community (Karst et al. 2023). However, using a metabarcoding approach, it has become possible to obtain a large amount of data describing fungal species present in soil communities (Tedersoo et al. 2010), which allows for comparative studies on fungal species composition. Previous metabarcoding studies have shown that fungal diversity depends on multiple factors, such as soil properties, elevation and humidity (Veresoglou et al. 2013; Sun et al. 2016; Zhang et al. 2021; Perez-Lamarque et al. 2022; Luo et al. 2023; Tedersoo et al. 2024). The composition and structure of RAF communities are highly influenced by local abiotic and biotic conditions (Alzarhani et al. 2019). However, the extent to which inter-species variation contributes to RAF community dynamics in naturally homogeneous environments remains uncertain. For instance, in an oak forest, roots of different plant species hosted distinct mycorrhizal and endophytic fungal communities, despite growing in proximity (Toju et al. 2013). In a tropical forest, Schroeder et al. (2019) found that the RAF community was more strongly associated with plant phylogeny than with spatial distance, with a stronger relationship observed for pathogens than for mutualists. At the same time, a metabarcoding study of different tropical communities revealed frequent fungal sharing between phylogenetically distant plant lineages at a fine scale, with significant variation in the level of specialization across different fungal lineages (Perez-Lamarque et al. 2022). The authors proposed that interaction of plants with fungi is determined by the intrinsic properties of each fungal lineage, but not by environmental conditions.

Within this study, we explore fine-scale variation in RAF communities among three cohabiting vascular plant species in a hemiboreal forest in Lithuania, dominated by *Pinus sylvestris* L. (Scots pine, Pinaceae). Alongside *P. sylvestris*, we studied the fine roots of two phylogenetically distant vascular plant species: the angiosperm *Vaccinium myrtillus* L. (European blueberry, Ericaceae) and the lycophyte *Diphasiastrum complanatum* (L.) Holub (northern groundcedar, Lycopodiaceae). Like many boreal forest trees, *P. sylvestris* is an obligate mutualist that relies highly on association with ECM fungi. As a member of the Ericaceae family, *V. myrtillus* is expected to form an obligate partnership with ERM fungi (Smith and Read 2008). The species composition of mycorrhizal fungi associated with *P. sylvestris* and *V. myrtillus* is relatively well studied (e.g., Rudawska et al. 2018; Sietiö et al. 2018; Olchowik et al. 2021; Vohník 2020; Bzdyk et al. 2022). However, there is limited knowledge about the fungal partners of lycophytes, the most basal extant lineage of vascular plants, particularly *D. complanatum* (Wikström and Kenrick 2001; Rimington et al. 2020). Gametophytes of lycophytes are often achlorophyllous and fully dependent on their symbiotic AM fungi for nutrition (and this is the case for *D. complanatum*; Horn et al. 2013), while the sporophytes may have a low dependence on their mycorrhizal partners when mature (Rimington et al. 2020). Lycophytes grow in boreal and hemiboreal forests alongside other vascular plants that evolved much later. The extent to which early and late diverging phylogenetically distant plant lineages share fungal partners remains a subject of ongoing debate (Rimington et al. 2018).

In this study, we conducted a metabarcoding analysis in combination with a light microscopic investigation to check two hypotheses: (i) each of the studied plant species hosts a distinct fungal species composition, with a prevalence of ECM fungi for *P. sylvestris*, ERM fungi for *V. myrtillus*, and AM fungi for *D. complanatum* sporophytes; (ii) the fungal communities associated with *V. myrtillus* fine roots exhibit the lowest species diversity due to the highest host specificity of ERM fungi.

## 2. Material and Methods

### 2.1. Study area

Our study area is located in Dzūkija National Park of Lithuania, with the investigated forest located on the north and south banks of the Merkys River (Fig. 1, Suppl. Tab. 1). The soils in this area have developed on non-carbonaceous quartz under sandy glacial plains, exhibiting podzolic characteristics with a mild podzolization, and are notably dry with severe deficiency in nutrients (Gulbinas and Samuila 2002). The annual gross solar irradiance in the study area is approximately 3300–3400 MJ/m^2^, the soil surface albedo is lower than 17.5%. The mean annual temperature is +6–7 °C, with an absolute maximum of +35.6 °C and an absolute minimum of −35.9 °C, and the temperature remains above 0 °C for 252 days a year. Mean annual precipitation is around 650 mm (National Land Service under the Ministry of Agriculture of the Republic of Lithuania 2016). The predominant tree species in the studied forest is Scots pine (*Pinus sylvestris*) (Augustaitis and Bytnerowicz 2008). According to the forest cadastre data (State Forest Service under the Ministry of Environment 2023), the age of the forest on the study area ranged from 76 to 126 years (average ± sd: 107 ± 15 years) and the height varied from 20.7 to 28.1 m (average ± sd: 26.0 ± 1.8 m) (Suppl. Tab. 1).

**Fig. 1.**
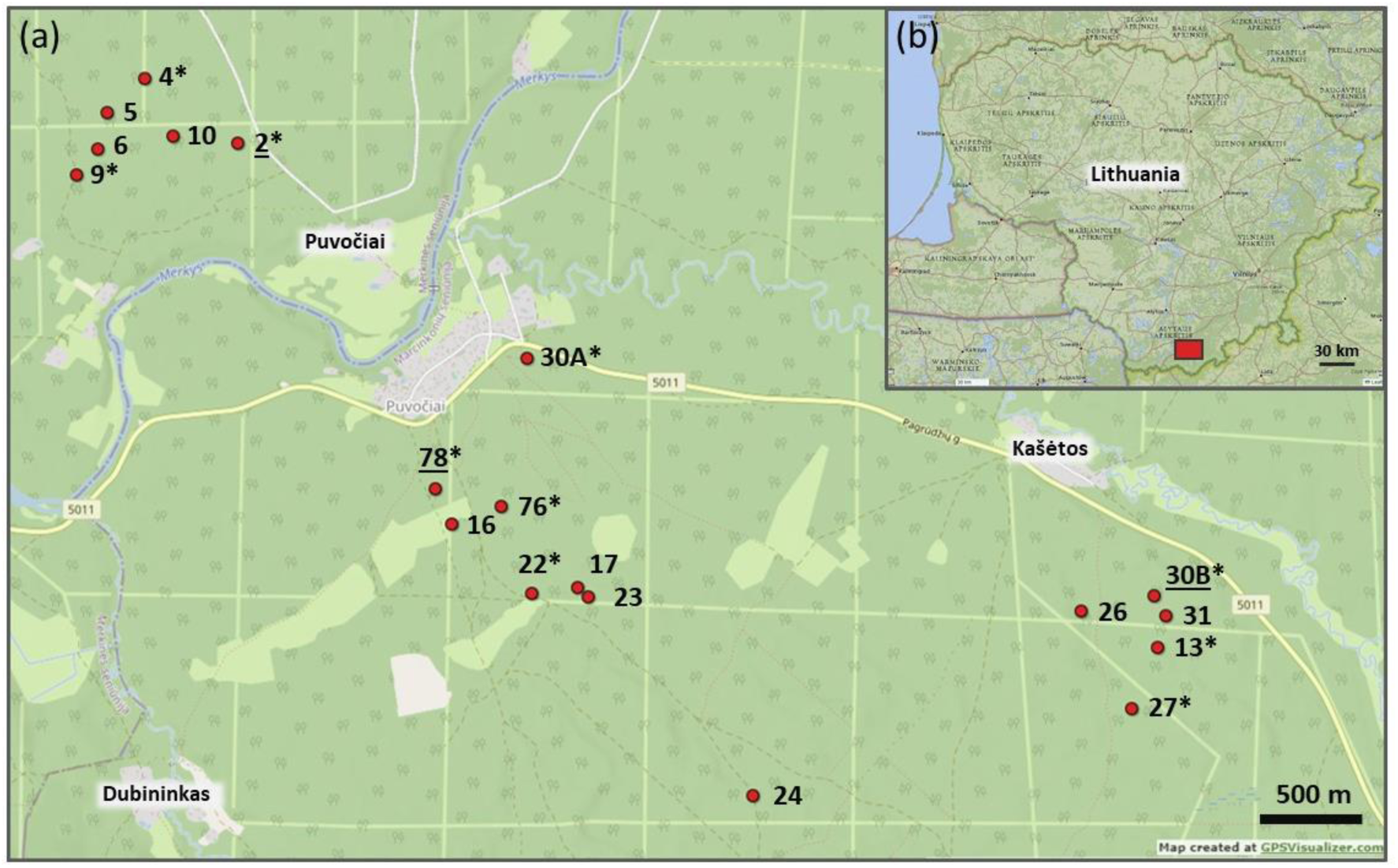
(a) Location of 19 plots (red circles) analysed via metabarcoding. Each plot was examined for fungi associated with soil and fine roots of *Pinus sylvestris*, *Vaccinium myrtillus*, and *Diphasiastrum complanatum* sporophytes. Labels with asterisks represent ten plots where fine roots of *D. complanatum* sporophytes were additionally analysed via microscopy; underscored labels indicate three plots where fine roots of all three plant species were investigated. (b) Location of the sampling region in southern Lithuania (red rectangle).

### 2.2. Sampling design

For metabarcoding analysis, samples of plant material and soil were collected in August 2021. Samples were collected from 19 plots (5 x 5 m each), where *Pinus sylvestris* L. co-occurred with *Vaccinium myrtillus* L. and *Diphasiastrum complanatum* (L.) Holub (Suppl. Tab. 1). In each plot, three samples of fine roots (5-10 cm length) of *P. sylvestris, V. myrtillus,* and *D. complanatum* sporophytes were collected from three different ramets/trees per species (Fig. 2). Additionally, soil samples were analysed to identify free-living or transient fungi and determine the "background" trophic modes characteristic of the local soil type: one teaspoon of soil was taken from three spots at a depth of 10 cm from the surface, and all visible root fragments were sorted out. The collection spoon was pre-treated with soil from the plot before sampling. Root samples were cleaned of large substrate fragments through gentle shaking and then washed by immersion in distilled water. Samples of the same plant species or soil from one plot were combined, resulting in a total of 76 samples. All samples were air-dried at room temperature (RT) and sealed in plastic zip-lock bags.

**Fig. 2.**
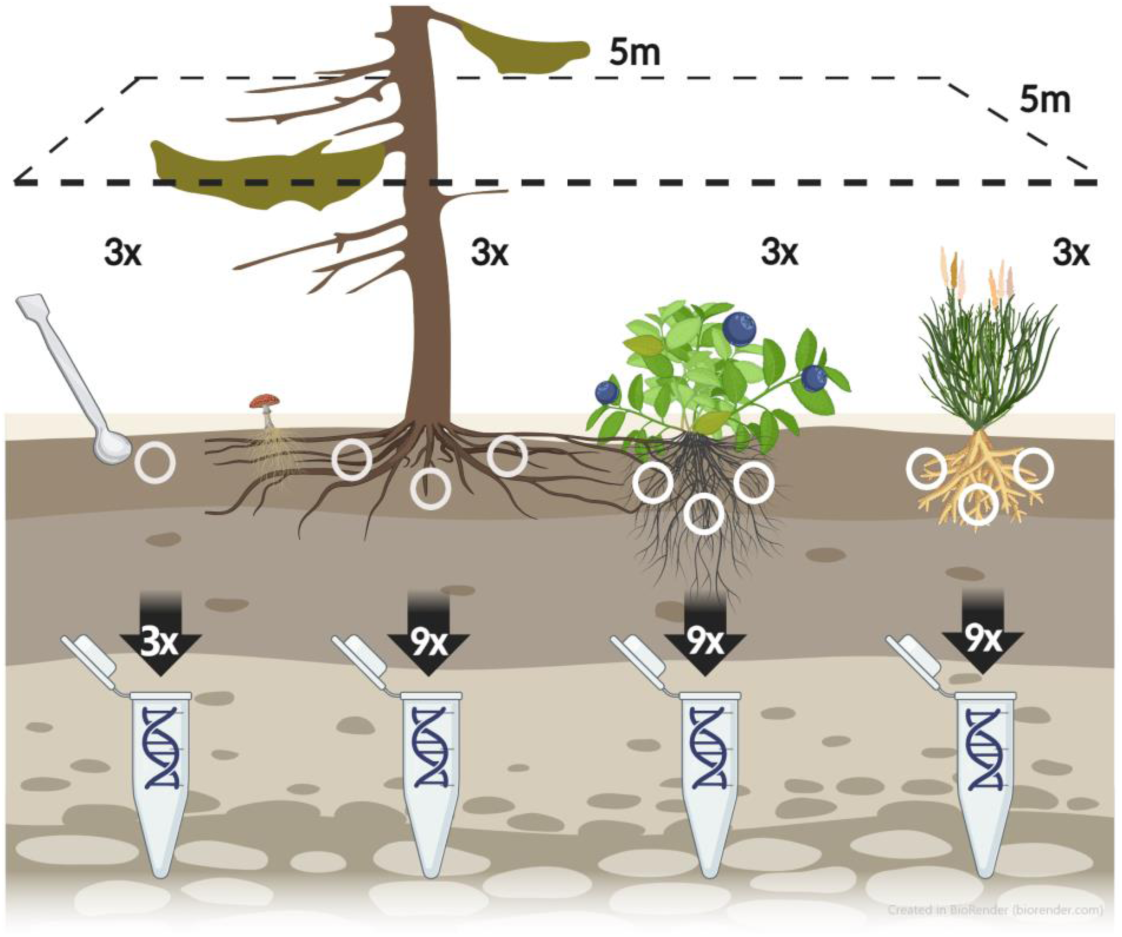
Collection scheme of samples for metabarcoding analysis of the soil and fine roots of *Pinus sylvestris*, *Vaccinium myrtillus*, and *Diphasiastrum complanatum* in each studied plot.

To characterize the vegetation of each plot, the coverage percentage of five strata, as well as of each plant and lichen species, was recorded. Sporophytes of *D. complanatum* often grow radially outwards, forming circle-shaped colonies known as "fairy rings" (Spalding et al. 1975). These colonies expand at a rate of 20 to 50 cm per year, with the central portions dying off. The diameter of these rings can be indicative of the colony’s age (Oinonen 1967), although re-invasion of the centre is possible. In our study, the diameter of the "fairy rings" ranged from 2 to 25 m (average ± SD: 10 ± 6 m; Suppl. Tab. 1), suggesting the age of the sporophytes could vary from 4 to 125 years.

For the microscopic analysis, additional sampling was conducted in August 2023, at ten previously investigated plots on both sides of the Merkys river (Fig. 2). Soil samples and fine roots of *D. complanatum* sporophytes were collected using the same sampling scheme, as for the metabarcoding analysis. Additionally, as positive standards for the preparation and staining, fine roots of *P. sylvestris* and *V. myrtillus* were sampled in three of these plots (“2”, “78”, ”30B”). The root material was preserved in 30% ethanol and stored at 6-8°C for three weeks. Soil samples were air-dried at RT and stored in paper bags.

The pH of each soil sample was measured according to DIN ISO 10390, after incubating 5 g of the soil in 25 mL of 0.01M CaCl_2_ overnight, with pH/mV pocket meter WTW pH330i and electrode WTW SenTix41 (both Xylem Analytics, Weilheim i. OB, Germany).

### 2.3. Metabarcoding library preparation

To avoid the batch effect, samples from a single plot were processed jointly during DNA extraction and amplification. Throughout the library preparation, the samples were quasi-randomly distributed across the plates.

For DNA extraction from plant material, 40 fragments of fine roots with ca. 2 mm length were selected from a single sample and pooled, resulting in an average of 15 mg of dry material. In cases where visible ECM root tips were present, they were preferentially chosen. Regarding soil samples, 300 mg of dry material was utilized. The DNA extraction procedure employed the NucleoSpin Soil extraction kit (Macherey-Nagel, Düren, Germany), following the manufacturer’s protocol with specific adjustments: root samples were dry homogenized in 2 mL tubes with a 5 mm steel ball, while soil samples were homogenized in provided tubes with added lysis buffers (SL and SX). The BeadBlaster 24TM rotor-stator homogenizer (Benchmark, Almelo, Netherlands) was employed for two cycles at 4.0 rpm for 1 min, with a 10 s break. The supernatant was transferred to a new tube before the precipitation of contaminants (step 4), and DNA was eluted in 50 μl of Elution buffer. In each of the four DNA extractions, a negative control of extraction was included. DNA extracts were quantified using a NanoDrop Lite Spectrophotometer (Thermo Scientific, Waltham, USA).

The amplification of the ITS2 region as a barcode was conducted using primers gITS7F (Ihnmark et al. 2012) and ITS4ngsR (Tedersoo et al. 2014), each incorporating Illumina overhang adapters (underlined): 5’-<u>TCGTCGGCAGCGTCAGATGTGTATAAGAGACAG</u>GTGARTCATCGARTCTTTG-3’ and 5’-<u>GTCTCGTGGGCTCGGAGATGTGTATAAGAGACAG</u>TTCCTSCGCTTATTGATATGC-3’, respectively.

The PCR was carried out in a 25 µl reaction mix comprising 12.5 µl of Taq 2x Master Mix RED with 1.5mM MgCl_2_ (Amplicon, Odense, Denmark), 0.2 µM of each primer, and 70 ng of DNA. The PCR protocol included the following steps: (i) 5 min at 94°C; (ii) 25 cycles of 30 s at 94°C, 30 s at 48°C, and 30 s at 72°C; (iii) 7 min at 72°C. A non-template negative control (NTC) was included for each of the two PCR runs. All samples (n = 76) and both types of negative controls (n = 5) were amplified; additionally, for all four samples from plot “2,” three technical replicates were utilized (n = 8). One NTC for NGS sequencing was also included, giving 90 libraries in total. Resulted PCR products were examined in 1.5% agarose gel with RotiSafe stain. All amplicon libraries were barcoded using the Illumina MiSeq v3 reagent kit for 2 × 300 bp sequencing, following the manufacturer’s protocol. After pooling, the libraries were sequenced in a single run at the Section of Microbiology (Copenhagen University, Denmark) where the sequences were also de-multiplexed, and the barcodes were removed.

### 2.4. Microscopic analysis

The preparation of root samples for microscopic analysis was conducted through Trypan blue staining following the modified protocol of Chabaud et al. (2006): (i) boiling at 90°C in 15 ml of 10% KOH (w/v) for 150 minutes; (ii) rinsing roots in tap water; (iii) bleaching in 15 ml of 10% H_2_0_2_ (v/v) with a few drops (0.2-0.3 ml) of 25% NH_3_ (v/v) for 2 hours; (iv) rinsing in tap water; (v) acidification of roots in 80% lactic acid for 1 hour; (vi) boiling at 90°C in 15 ml of 0.2% Trypan blue solution in lactoglycerol for 3 minutes. Stained roots were preserved in lactoglycerol at 4-6°C. Subsequent mycorrhiza detection was conducted using a Leica DM2500 light microscope with FLEXACAM C1 camera (both Leica Camera, Wetzlar, Germany) and the associated program Leica Application Suite v4.8.0.

Since *D. complanatum*, the least studied species in this context, was the primary focus of our microscopic investigation, we examined a total of 300 fine root fragments from its sporophytes (Suppl. Fig. 1c). Additionally, 90 fragments each from *P. sylvestris* and *V. myrtillus* were examined, primarily as positive controls for sample preparation and staining. For each sample, 30 fragments of fine roots, each minimum 1 cm long, were analysed for the presence/absence of ECM, AM, or ERM via presence of arbuscles, hyphal mantles, or hyphal coils. To identify ECM, we looked for fine root tips that exhibit swollen, chromatically shiny tips, typically dichotomously branched in *Pinus*. The presence of arbuscules or hyphal coils within root cells indicates AM or ERM, respectively.

### 2.5. Data analysis

Nextera adapters, as well as N and low-quality leading and trailing bases (<3, Phred-33), were trimmed using Trimmomatic v0.39 (Bolger et al. 2014). Quality of the raw and resulting sequences was checked by running FastQC v0.11.5 and MultiQC v1.19 tools (Andrews 2010; Ewels et al. 2016). Demultiplexed and trimmed reads are available on NCBI (PRJNA1185013).

Obtained sequences were processed using the VSEARCH v2.15 Python package (Rognes et al. 2016). The following steps were undertaken: merging of forward and reverse reads (*fastq_minovlen* = 150, *fastq_maxdiffs* = 15), quality filtering of reads (*fastq_maxee* = 0.5, *fastq_minlen* = 250, *fastq_maxns* = 0), dereplication of reads across the samples and removal of singletons (*minuniquesize* = 10), pooling of the samples, and eliminating reads with putative PCR errors (denoising; *fastq_eeout* parameter). The VSEARCH v2.15 package was also utilized for the identification and removal of chimera sequences using the UCHIME algorithm, applying both *de novo* and reference-based detections (Edgar et al. 2011). All resulting sequences with a similarity lower than 97% were clustered to the different Operational Taxonomic Units (OTUs) using “greedy” clustering in the function *cluster-size*. Each OTU was taxonomically assigned by comparing it to the UNITE v9.0 database with a 96% identity threshold (Abarenkov et al. 2023). As a result, multiple OTUs could be assigned to the same fungal taxa. The assignment of the bottom ten OTUs with the lowest match values was additionally verified by blasting them against the GenBank database (megablast, nr/nt). For ecological characterization of revealed genera and species, FUNGuild v1.1 database was implemented to assign them to guilds and trophic modes (Nguyen et al. 2016). If a taxon was assigned to multiple guilds, including ECM, it was categorized as “ectomycorrhizal”. Taxa assigned to multiple guilds exclusively within the saprotrophic category, including pathogenic and endophytic, were categorized as “saprotrophic”. Guilds and trophic modes were assigned to taxa only when confidence levels were “highly probable” or “probable”.

Statistical evaluation and visualisation of results were conducted in R 4.4.2 using RStudio 2023.12.1 (RCore Team 2024; Posit Team 2024).

Prior to conducting the data analysis, OTUs potentially associated with contamination were excluded. This process involved removing OTUs with total counts from NTCs that exceeded a half of the normalized total sample counts, obtained by dividing the total sample counts by the ratio of the number of samples to the number of NTCs (84/6). NTCs were then removed from the dataset. Following this treatment, the counts underwent normalization for sampling depth through a variance stabilizing transformation implemented in the R package *DESeq2* v1.46 (Love et al. 2014). After this step, each count value within a sample underwent subtraction by the lowest count value in that sample, resulting in zero for absent counts.

To visualise variations in taxonomic composition among sample types, a Non-metric Multidimensional Scaling (NMDS) with Bray distances, as well Principal Coordinate Analysis (PCoA) with Euclidean distances (with and without technical replicates) were performed utilizing the R package *phyloseq* v1.50 (McMurdie and Holmes 2014). Three types of datasets were used for NMDS: (1) one containing all revealed 278 OTUs, (2) one containing only OTUs assigned to ectomycorrhizal functional guilds of fungi (86 OTUs), (3) and one with only OTUs of saprotrophic fungi (108 OTUs). The outcomes were graphically represented using the R packages *ggplot2* v3.5.1 and *ggrepel* v0.9.6 (Wickham 2016; Slowikowski 2024). In the subsequent data analysis steps, technical replicates were excluded.

Taxonomic relative abundance per sample type was visualized using the R packages *microeco* (v1.11.0) and *ggplot2* (Liu et al. 2021). The statistical significance between groups of samples at each taxonomic rank was determined using the Wilcoxon Rank Sum test, with the false discovery rate *p*-value correction for multiple comparisons (Benjamini and Hochberg 1995; Haynes 2013) using the R package *metacoder* v0.3.7 (Rice et al. 2000). Four types of pairwise comparisons of relative abundances were conducted: (1) between different sample types, (2) between *P. sylvestris* samples categorized by high and low tree age, (3) between *P. sylvestris* samples categorized by high and low tree height, and (4) between *D. complanatum* samples categorized by large and small colony diameters. Samples were classified into high or low groups based on whether their respective values were above or below the corresponding average. To achieve this, normalized counts of OTUs across all ranks of every taxon were aggregated within a sample type. Only taxa represented by at least two OTUs were included in the analysis to simplify visualization and interpretation of the results. In the case of comparison of the sample type, results were depicted on a heat tree.

To visualize OTU diversity within each sample type, sample-based rarefaction and extrapolation curves were generated for Hill numbers (OTU richness, Shannon, and Simpson diversities) using the R package *iNEXT* v.3.0.1 (Hsieh et al. 2022). Additionally, this package was employed to construct a sample completeness curve to assess the adequacy of the sample size. A Venn diagram was built to show the number of unique and shared OTUs across sample types using the online tool *jvenn* (Bardou et al. 2014).

The abundance of trophic modes and guilds among sample types was statistically tested. Prior to analysis, the dataset was assessed for normality using the Shapiro-Wilk test and for homogeneity of variances using Bartlett’s test. Normally distributed data with homogeneous variances were analysed using one-way analysis of variance (ANOVA) followed by Tukey’s honest significant difference (HSD) test (Tukey 1949), while data not meeting these assumptions were analyzed using the Kruskal-Wallis test with Dunn’s post-hoc test, implemented using the R package *rstatix* v0.7.2 (Kassambara 2023). For both tests, Bonferroni correction was applied to adjust *p*-values for multiple comparisons at a significance level of α = 0.05.

A MultiQC report on the raw and trimmed reads, along with raw and normalized read counts, taxonomic assignment of OTUs, and scripts in bash (Trimmomatic, VSEARCH, and FUNGuild steps) and R are available on GitHub: https://github.com/kuprinak/lithuania-metabar.

## 3. Results

### 3.1. Plot characteristics

All sampling plots had acidic soils, with pH varying from 2.9 to 4.0 across two sampling years (average pH ± sd: 3.3 ± 0.18 in 2021, 3.4 ± 0.25 in 2023, Suppl. Tab. 1).

The vegetation survey conducted on sampling plots revealed *P. sylvestris* as the dominant tree species with 28.0% surface coverage, followed by *Betula pendula* Roth (6.1% coverage) (Suppl. Tab. 2). The dominant shrubs and subshrubs observed were *Juniperus communis* L. with 9.3% coverage, *Vaccinium vitis-idaea* L. with 6.6%, *D. complanatum* with 6.3%, and *Vaccinium myrtillus* with 4.6%. Typically, each plot contained a single individual of *D. complanatum* consisting of multiple shoots connected by subterranean rhizomes. Previous analysis revealed that the individuals genetically differed between plots (Schnittler et al. 2019). Interestingly, the tree and shrub strata comprised both typical AM plants, such as *Prunus serotina* Ehrh*., Juniperus communis, Sorbus aucuparia* L., as well as ECM plants like *Pinus sylvestris*, *Picea abies* (L.) Karst., *B. pendula*, and *Quercus robur* L. (Akhmetzhanova et al. 2012). Mosses and lichens collectively covered an average of 75.4% of the ground surface, with the bryophytes *Hylocomium splendens* (Hedw.) Schimp. and *Pleurozium schreberi* (Brid.) Mitt. being most prevalent.

### 3.1. Metabarcoding analysis

#### 3.1.2. DNA library

Sample sequencing produced a total of 13,938,499 raw read pairs, with an average of 165,934 read pairs per sample and standard deviations of 29,378 read pairs across samples and 6,100 read pairs across sample types. For the NTCs, 31,158 read pairs were produced, averaging 5,193 read pairs per NTC. After processing reads with Trimmomatic, 10,128,361 read pairs remained, with an average of 120,500 read pairs per sample. The average sequence length of the libraries ranged from 220 to 273 bases, with a mean of 252 bases. All sample libraries successfully passed quality checks, including per-base N content and adaptor content assessments, as evaluated by FastQC and MultiQC tools.

After processing the data with the VSEARCH package, 283 distinct OTUs were initially identified. Four OTUs were excluded due to potential contamination, and one was excluded because no counts remained after NTC removal, reducing the total to 278 OTUs.

Taxonomic assignment successfully classified 99.6% of these OTUs to the phylum, 95.1% to the class, 87,6% to the order, 80.9% to the family, 73.5% to the genus, and 47.0% to the species level. In total, nine phyla, 21 class, 37 orders, 62 families, 77 genera, and 132 species of fungi were revealed (Suppl. Fig. 2). The most abundant phylum was shown to be Basidiomycota with Agaricomycetes as the most abundant class (Suppl. Fig. 3). The most species rich genera were *Cortinarius*, *Russula*, and *Mortierella* with 15, nine and eight species, respectively.

#### 3.1.3. Species composition among sample types

Two distinct clusters were identified through NMDS analysis: (1) soil samples grouped with *D. complanatum* and (2) *P. sylvestris* with *V. myrtillus* (Fig. 3a). PCoA, however, revealed three clusters: (1) soil samples, (2) *P. sylvestris* and *V. myrtillus*, and (3) *D. complanatum* positioned in between (Fig. 3d). Technical replicates for each sample were tightly clustered (Suppl. Fig. 4).

**Fig. 3.**
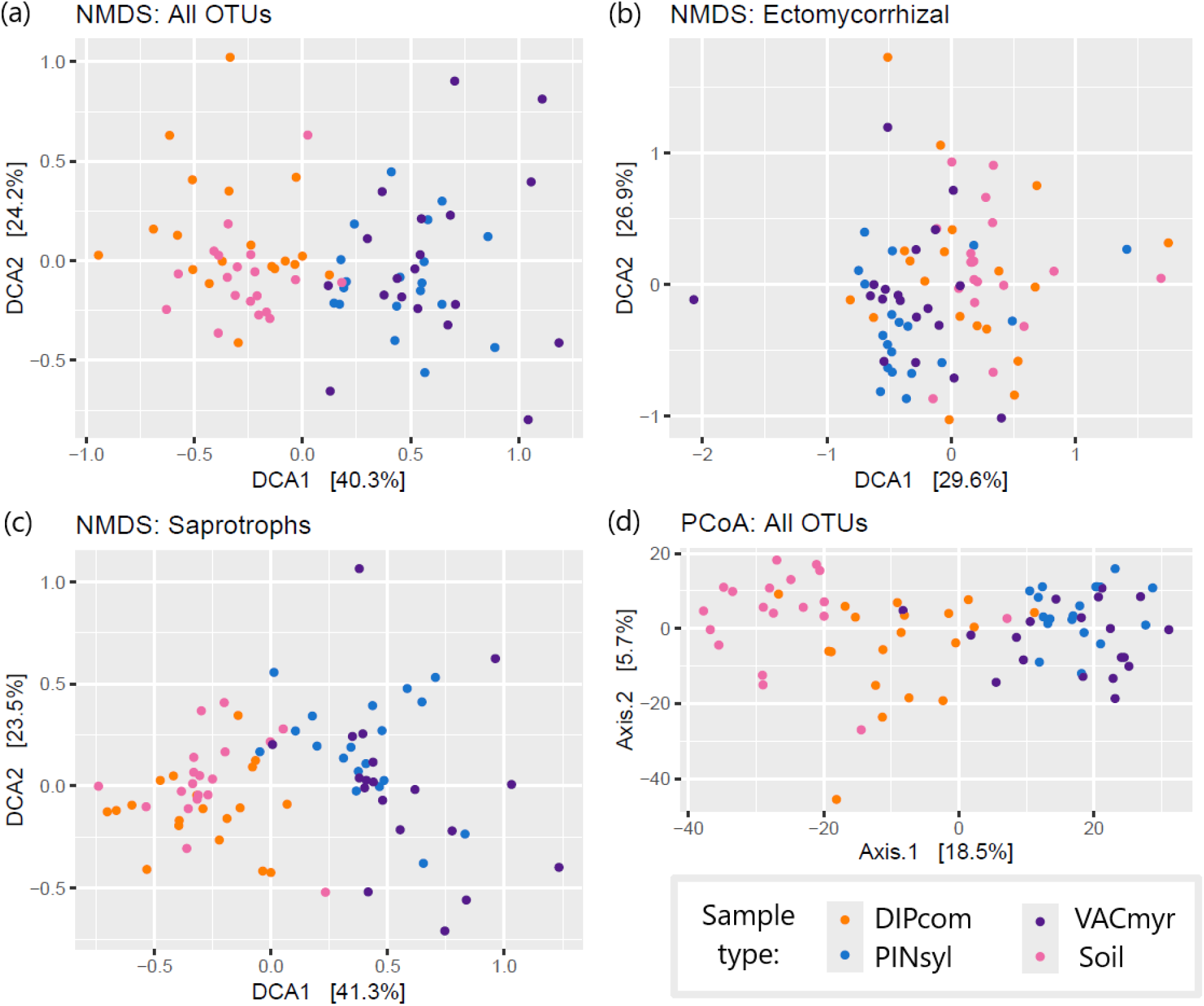
Ordination diagrams generated from Bray distances via Non-metric Multidimensional Scaling (NMDS) for all (a) 278 OTUs, (b) 86 OTUs of ectomycorrhizal fungi, and (c) 108 OTUs of saprotrophic fungi. (d) Ordination diagram generated from Euclidean distances via Principal Coordinate Analysis (PCoA) for all 278 OTUs. Distances were calculated between normalized counts of OTUs found in soil, *Diphasiastrum complanatum* sporophytes (DIPcom), *Pinus sylvestris* (PINsyl), and *Vaccinium myrtillus* (VACmyr) fine root samples collected in 19 plots (83 samples in total).

The number of observed OTUs varies among sample types, with the highest number (256 OTUs) found in soil samples and the lowest (201 OTUs) in *P. sylvestris* samples (Suppl. Fig. 5b). More than half of all OTUs (182 OTUs, 66%) were present in all four sample types, while only soil samples contained OTUs that were not found in any other sample type (26 OTUs, 9%) (Suppl. Fig. 5a).

Significant differences in relative abundancies across the sample types were shown for multiple taxa (Figs. 4a-g). The highest number of significant differences was found between soil samples and those from *P. sylvestris* and *V. myrtillus*: soil had predominance across six Phyla; only the genus *Luellia* was more abundant in *P. sylvestris* and *V. myrtillus*, and the genus *Mycena* was dominant in *V. myrtillus* samples (Figs. 4e, 4f). Samples of *D. complanatum* showed only genus *Mucor* as a more prevalent taxon in comparison with the soil samples (Fig. 4g). At the same time, several taxa were dominant in *D. complanatum* samples in comparison with *P. sylvestris* and *V. myrtillus* samples: the phyla Mucoromycota (classes Mucoromycetes and Umbelopsidomycetes), Ascomycota (class Sordariomycetes), Rozellomycota, and Basidiomycota (classes Tremellomycetes, Geminibasidiomycetes, and the genera *Amanita* and *Ceratobasidium* from the class Agaricomycetes) (Figs. 4b, 4c). Less taxa with significant differences were found between samples of *P. sylvestris* and *V. myrtillus*: *P. sylvestris* had more reads of the genus *Russula*, while *V. myrtillus* had more abundant reads of the family Serendipitaceae (Sebacinales), the order Auriculariales, and the genus *Trechispora* (Trechisporales) (Fig. 4d).

**Fig. 4.**
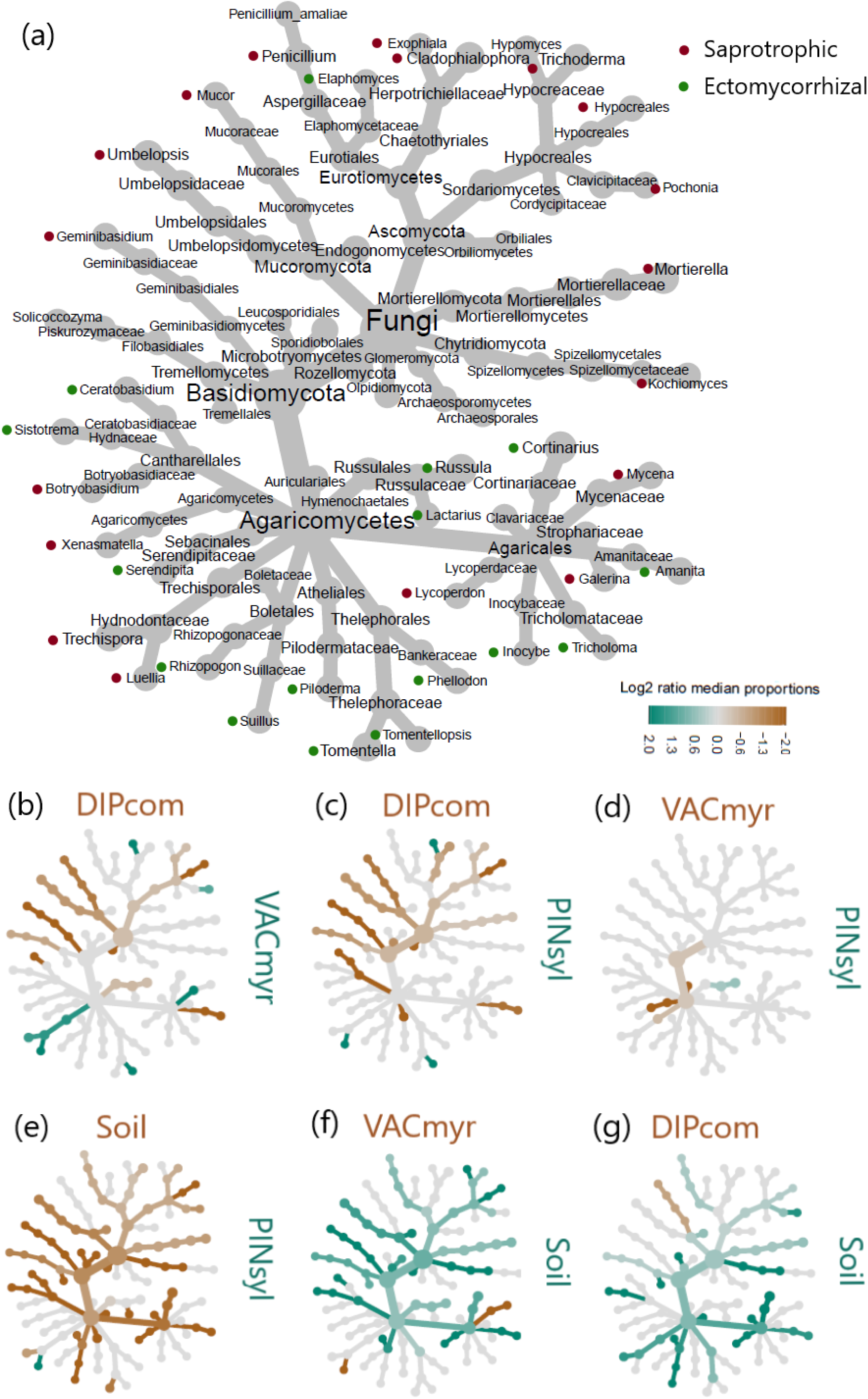
(a) Tree of fungal taxa identified during metabarcoding analysis of four sample types (only taxa with OTUs number ≥ 2 included): soil, fine roots of *Diphasiastrum complanatum* sporophytes (DIPcom), *Vaccinium myrtillus* (VACmyr), and *Pinus sylvestris* (PINsyl) in 19 studied plots (76 samples in total). The genus labels are colour-coded according to guilds defined using the FUNGuild v1.1 database. (b - g) Heat trees illustrating the comparison of abundance of normalised counts of OTUs assigned to the corresponding taxa among sample types. The colour of each taxon node represents the log-2 ratio of median proportions of reads observed at sample type determined by a Wilcox rank-sum test with a false-discovery rate correction for multiple comparisons. Taxa coloured green have significantly higher number of reads in the sample type shown in the row and the taxa coloured brown are prevalent in the sample type shown in the column.

No significant difference in taxa relative abundancies was found between samples from plots with high and low tree age or height for both types of datasets (all types of samples and only *P. sylvestris* samples), as well as between *D. complanatum* samples with “big” and “small” colony sizes.

#### 3.1.4. Relative OTU diversity among sample types

All calculated diversity indices consistently indicated a similar trend: soil samples exhibited the highest values of Hill numbers, followed by *D. complanatum* and *V. myrtillus* samples, and the lowest values recorded for *P. sylvestris* samples. (Figs. 5a-c). The sample completeness curves reached the plateau shape for each sample type, indicating an adequacy of the sample number for analysing species diversity (Suppl. Fig. 6). The pattern remained consistent when only the 86 OTUs assigned to ECM fungi were considered (Suppl. Fig. 7).

**Fig. 5.**
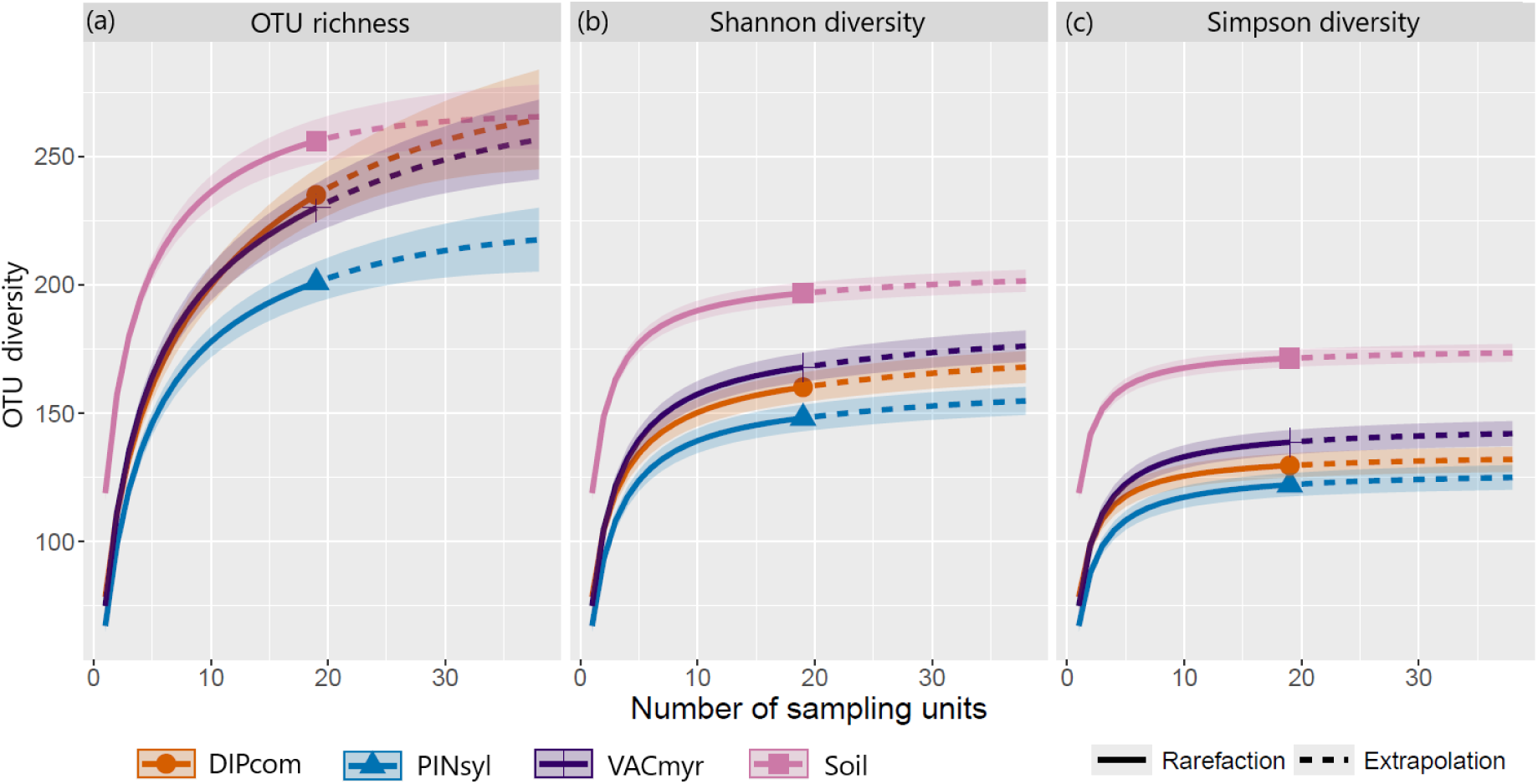
Rarefaction and extrapolation curves with 95% confidence intervals of three Hill numbers describing OTU diversity: (a) OTU richness, (b) Shannon diversity, (c) Simpson diversity. The numbers are calculated using normalized counts of 278 OTUs for soil or fine roots of *Diphasiastrum complanatum* sporophytes (DIPcom), *Pinus sylvestris* (PINsyl) and *Vaccinium myrtillus* (VAC myr) from 19 studied plots (76 samples in total)

#### 3.1.5. Trophic modes

Of the 278 OTUs, FUNGuild assigned 218 OTUs to their functional guilds, with 86 categorized as ECM and 108 as saprotrophic fungi. The most abundant ECM genera were *Russula*, *Cortinarius*, *Tomentella*, and *Amanita*, while the most abundant saprotrophic genera included *Umbelopsis*, *Mortierella*, *Penicillium*, *Cladophialophora*, and *Trichoderma* (Suppl. Fig. 3e).

Plots of NMDS constructed exclusively for ECM OTUs separated the samples into two distinct clusters: (1) soil samples grouped with *D. complanatum* and (2) *P. sylvestris* grouped with *V. myrtillus* (Fig. 3b). In contrast, clustering was less distinct for the saprotrophic functional guilds (Fig. 3c).

The relative abundance of fungal trophic modes and functional guilds differed between sample types (Figs. 6a,b). Compared to the other sample types, roots of *D. complanatum* sporophytes showed significantly lower abundance of ECM fungi and symbiotrophs in comparison to *P. sylvestris* (adjusted *p*-values from the Kruskall-Wallis test: *p* = 0.0105 and *p* = 0.0007, respectively). Samples of *D. complanatum* also comprised a significantly higher abundance of OTUs with saprotrophic modes than *P. sylvestris* (adjusted *p*-value from the Tukey HSD test: *p* = 0.0025). Soil samples contained more saprotrophs than *D. complanatum, P. sylvestris* or *V. myrtillus* (adjusted *p*-values from the Tukey HSD test: 0.0274, 0.0205, 0.0275, respectively).

**Fig. 6.**
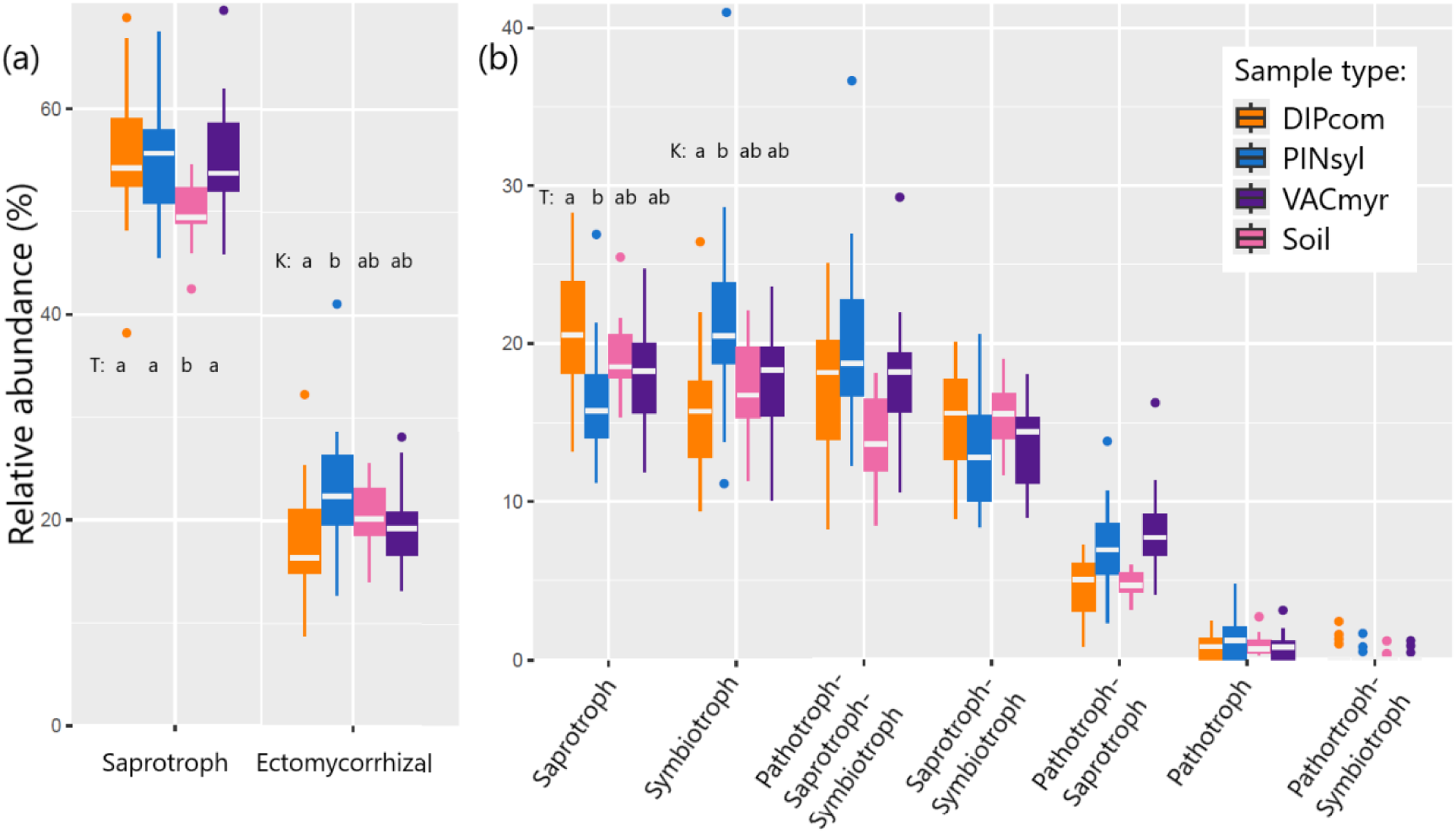
Relative abundance of normalized counts of 278 OTUs assigned to (a) functional guilds and (b) fungal trophic modes using FUNGuild v1.1 database for soil of or fine roots of *Diphasiastrum complanatum* sporophytes (DIPcom), *Pinus sylvestris* (PINsyl) and *Vaccinium myrtillus* (VAC myr) from 19 studied plots (76 samples in total). Each bar represents one sample (n = 76). Boxplots show differences between sample types for the two most abundant (b) trophic modes and (c) guilds. Samples of the boxes with the same letter code are not significantly different (T – Tukey Post-hoc test, K - Kruskal-Wallis test with the Dunn post-hoc test).

### 3.3. Microscopic analysis

A distinctive feature of *D. complanatum* fine roots was a dense cover of unicellular root hairs, visible to the naked eye (Suppl. Figs. 1b, 1c). Investigation of all 300 fine root fragments from *D. complanatum* sporophytes revealed an absence of arbuscles, hyphal mantles, or hyphal coils characteristic to AM, ECM and ERM, respectively (Figs. 7a, 7b). Root cells primarily contained remnants of cell components, with unstructured hyphae occasionally covering them. In contrast, all examined fragments of *P. sylvestris* fine root samples exhibited mantles of ECM hyphae (Fig. 7c), and all hair root fragments from *V. myrtillus* contained cells with hyphal coils of ERM (Figs. 7d, 7e).

**Fig. 7.**
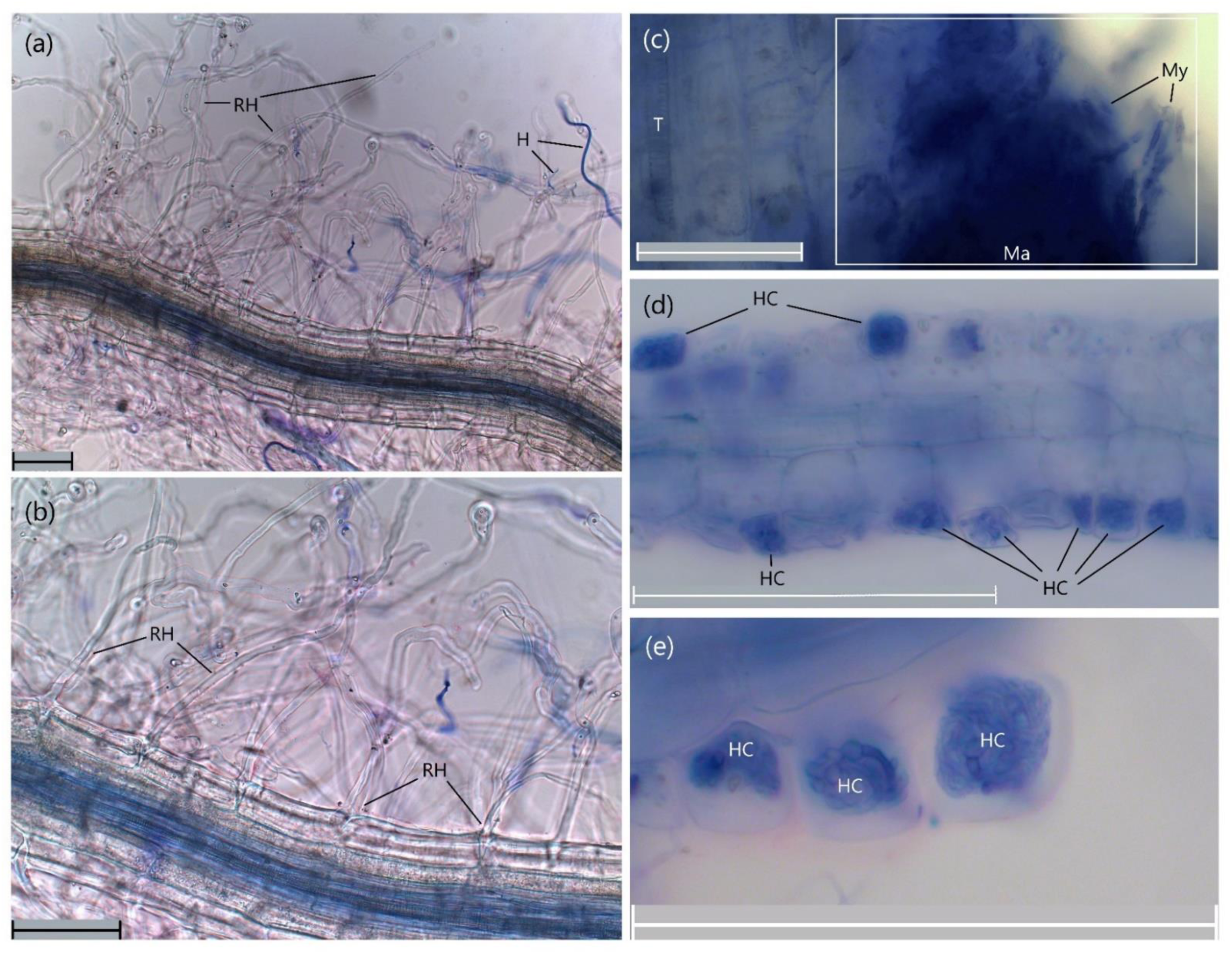
(a, b) Fine root of *Diphasiastrum complanatum* sporophyte; (c) fine root of *Pinus sylvestris*; (d, e) hair root of *Vaccinium myrtillus*. RH – root hair, H – hypha of fungi, Ma – mantel of hyphae, My – mycelium, HC – hyphae coils. Scale bar - 100 µm.

## 4. Discussion

### 4.1. Fungal community

In the studied hemiboreal Scots pine forest, a diverse fungal community is present, with Basidiomycota as the most prevalent phylum and saprotrophic fungi as the dominant trophic mode. In boreal forests, ECM is the predominant mycorrhizal type, playing a crucial role in nutrient cycling (Smith and Read 2008, Clemmensen et al. 2013; Kyaschenko et al. 2019). Previous studies have shown that ECM fungi are most abundant in soils of forests dominated by Pinaceae and can comprise about 34% of all fungal taxa (Tedersoo and Smith 2013; Tedersoo et al. 2012). In our study, only 20% of fungal taxa from the soil samples were assigned to the ECM functional guild, and almost 50% to saprotrophic fungi (Fig. 6a). However, functional overlap between these groups is possible: saprotrophic fungi can form facultative symbiotrophic relationships with plant roots, while some ECM fungi exhibit capabilities for facultative saprotrophy and decomposition (Koide et al. 2008; Bödeker et al. 2014; Smith et al. 2017). Additionally, the fungal community is a dynamic system which can change over time and especially after rapid environmental changes (Rudawska et al. 2018; Taniguchi et al. 2018; Olchowik et al. 2021). Various genera of ECM fungi are associated with different forest or host ages. For example, the genus *Russula* was shown to be more diverse and abundant in older *P. sylvestris* stands (above 50 years), while the genera *Lactarius* and *Suillus* were associated with younger stands (Dahlberg and Finlay 1999; Hutchison 1999; Brundrett 2002; Rudawska et al. 2018). Given the relatively old age of the trees in the studied plots (average 102 years), our results are comparable with data from an 87-year-old *P. sylvestris* forest in Poland, which also showed a predominance and highest species-richness of *Russula* and *Cortinarius* (Rudawska et al. 2018). A metabarcoding study of the boreal forest in Finland identified the ECM-forming Basidiomycetes *Cortinarius*, *Russula*, *Piloderma*, and *Tomentella* as the most abundant genera in the soil (Sun et al. 2016). Similarly, in our study, *Cortinarius*, *Russula*, and *Tomentella* were the most abundant ECM genera, with the exception of *Piloderma*, which ranked after *Tomentella* and *Amanita* (Suppl. Fig. 3e). Overall, however, saprotrophic *Umbelopsis* and *Mortierella* were the most common genera. *Cortinarius*, *Russula*, and *Mortierella* were also the most species-rich, likely representing the core fungal community in the studied forest.

Despite being most abundant in temperate grasslands and tropical rainforests, AM fungi have also been found in temperate forests, including those dominated by *P. sylvestris* (Vandenkoornhuyse et al. 2002; Onguene and Kuyper 2001; Öpik et al. 2003). However, our microscopic analysis did not reveal any signatures of AM colonisation in any type of samples. Tedersoo et al. (2016) reported that two AM genera, *Glomus* and *Endogone,* have a lower number of rDNA copies than the other studied fungi taxa, leading to low amplification of their ITS regions. In our study, three OTUs were assigned to the phylum Glomeromycota (Suppl. Fig. 2), but no significant difference in the relative abundance were found between sample types (Figs. 4a-g). Hovewer, the Mucoromycota was relatively abundant and included the species *Umbelopsis dimorpha, U. angularis*, *Mucor hiemalis, M. moelleri, M. sylvaticus,* and *Bifiguratus adelaidae*. The latter species is frequently sequenced from soil in northern temperate zones and may have a symbiotic function, having been detected in orchid and chestnut roots (James and Seifert 2017). *Umbelopsis dimorpha* was also described as an endopyte (Quin et al. 2018).

Despite previously reported difficulties in detecting closely related Mucoromycota and Mortierellomycota fungi using the ITS2 marker (Tedersoo et al. 2016; Perez-Lamarque 2022, 2023), these phyla were the third and fourth most abundant in our dataset, respectively (Suppl. Fig. 3a). This lower amplification bias among phyla can be attributed to using a different primer combination, gITS7F-ITS4ngsR, instead of ITS86F-ITS4. Different genetic markers and primer combinations can lead to biased detection of specific fungi in a sample due to varying primer specificities (Waud et al. 2014; Op De Beeck et al. 2014). Moreover, mycorrhizal root colonization can fluctuate seasonally and has a limited lifespan, potentially making it undetectable if not sampled at the right time (Majdi et al. 2001; Kemp et al. 2003; Mandyam and Jumpponen 2008; Meddad-Hamza et al. 2017). Therefore, it is important to be cautious when comparing the results of the studies utilizing different sampling and DNA library preparation techniques.

### 4.2. Species composition among sample types

Despite analysing samples of different species collected within an area of five square meters, this study found significant differences in fungi composition among sample types. Surprisingly, the smallest difference in the relative abundance of taxa was found between samples of *V. myrtillus* and *P. sylvestris* (Fig. 4d), two species known to form different types of mycorrhizal symbiosis.

#### 4.2.1. Diphasiastrum complanatum

It has long been known that Lycopodiaceae species rely upon mycorrhizae for successful development of both sporophytes and gametophytes (Bruchmann 1898; Whittier 1977; Schmid and Oberwinkler 1993; Winther and Friedman 2008, Horn et al. 2013). Surveys of mycorrhizal associations in lycophyte sporophytes often report not only AM, but also ERM, orchid mycorrhiza, and microsclerotia with Ascomycota and Basidiomycota (Spessard 1917; Freeberg 1962; Treu et al. 1996). However, our knowledge about lycophyte fungi interactions and fungal sharing remains scarce (Horn et al. 2013, Perez-Lamarque et al. 2023). Rimington *et al*. (2020) reviewed publications on the fungal symbiosis status of lycophytes and found that only six species showed Mucoromycotina, 53 had Glomeromycota and one species Basidiomycota. However, the authors pointed out that Mucoromycotina symbionts have likely been misidentified as Glomeromycota and the number of species with Glomeromycota could be lower.

The ordination analyses (NMDA and PCoA) revealed that the fungi associated with *D. complanatum* sporophyte roots are a distinct community and more closely related to the fungal community in soil samples than to that associated with *P. sylvestris* and *V. myrtillus* (Figs. 4a-d). Moreover, the Hill numbers describing OTU diversity of ECM fungi were more similar to the soil samples, than other sample types (Suppl. Figs. 8a, 8b, 8c). This similarity to the soil samples might be connected to the fragility and high surface area of the fine roots, which made them difficult to separate from soil organic particles during the cleaning process. Hovewer, the Hill numbers calculated with all OTUs were significantly lower for *D. complanatum,* than for the soil (Figs. 5a, 5b, 5c). This observation could be related to the different weight of the analyzed roots compared to the soil samples (15 mg vs. 300 mg), despite normalizing the amount of DNA used for amplification and library preparation. The sample coverage plot (Suppl. Fig. 6) showed that each sample type reached a plateau, but the soil samples exhibited significantly higher coverage than the root samples, merging only after 35 samples per type. Nonetheless, all root samples showed no significant differences in their coverage, supporting the robustness of the comparison.

Only the order Mucoromycetes was more abundant in *D. complanatum* samples than in the soil samples (Figs. 4a, 4g). However, in comparison with *P. sylvestris* and *V. myrtillus*, there was a predominance of such saprotrophic genera as *Solicoccozyma*, *Geminibasidium*, *Umbelopsis, Mucor, Trichoderma*, and *Hypocreales* (Figs. 6c, 6e). Only two ECM genera, *Ceratobasidium* and *Amanita*, were more abundant in *D. complanatum* samples compared to both *P. sylvestris* and *V. myrtillus*. However, soil samples also showed predominance of these ECM genera compared to *P. sylvestris* and *V. myrtillus* samples (Figs. 4e, 6f). It is not yet clear whether these fungi participate in any types of mycorrhizal partnership with *D. complanatum* and if so, these fungi behave rather opportunistic.

Perez-Lamarque *et al*. (2023) demonstrated that gametophytes and sporophyte roots for *Diphasiastrum tristachyum*, *D. oellgaardii*, *D. zeilleri,* and *Lycopodium clavatum* were colonized by a specific fungus from the clade Densosporaceae (Endogonales, Mucoromycotina). This fungus was abundant in the germinated spores suggesting that lycopod spores require this specific association for germination. For the roots of sporophytes, they found a dominance of Leotiomycetes, Archaeorhizomycetes, Endogonales and the parasitic genus *Spizellomyces* (Chytridiomycota). In our study, OTUS of Endogonomycetes and *Spizellomyces* were found, and their relative abundances in *D. complanatum* samples were not higher than in the other root samples (Figs. 4b, 4c) and were even lower than in the soil samples (Fig. 4g). This discrepancy between studies could be attributed to differences between studied biotopes: plants from the previous study were collected from artificially created subalpine heathland after the disturbance of a beech forest in the Hochfeld reserve, France. Perhaps another group of fungi plays a crucial role in the germination of *D. complanatum* spores in the pine forests in Lithuania and it has yet to be identified.

The surprising observation was that the multiple root hairs of *D. complanatum* sporophytes stayed intact after boiling the roots at 90°C in 10% KOH for 150 minutes (Figs. 7a, 7b). Similarly, root hairs of *Lycopodiella inundata* stayed intact after 20 min of boiling (Kowal et al. 2020). The cell walls of lycophyte root hairs may differ structurally and be more resilient than those of angiosperm plants, potentially due to the parallel evolution of lycophyte roots alongside other seed plants (Raven and Edwards 2001; Weng et al. 2008; Weng and Chapple 2010). Further research into the root hair chemistry and anatomy of lycophytes could provide deeper insights into this unique structure.

In our study, we did not find any microscopic evidence for the presence of AM, ECM, or ERM in the fine roots of *D. complanatum* sporophytes. Multiple septated hyphae could be observed around the root, which however did not form any mycorrhiza-like structures. Occasional vesicles were observed inside the fine root cells, resembling Mucoromycotina endophytes previously identified in the fine roots of *Lycopodiella inundata* sporophytes (Kowal et al. 2020). The presence of long and dense root hairs was a peculiar and distinguishing feature of all *D. complanatum* fine root samples, visible even to the naked eye (Fig. 9a, Suppl. Fig. 1). The root hairs closely resemble ’root clusters’ found in several non-mycorrhizal seed plants, which enhance phosphorus uptake from nutrient-poor soils (Lambers et al. 2006)—an adaptation that may have evolved independently in *D. complanatum*. We speculate that, unlike the obligatory mycorrhizal plants *P. sylvestris* and *V. myrtillus*, the lycophyte *D. complanatum* is less dependent on fungi partnership due to the increased surface resulting from numerous root hairs.

Only “adult” sporophytes, usually clones with numerous above-ground shoots, were investigated in this study, and no sporophyte that was still connected to its gametophyte was collected. The diameter of the “fairy rings” of the colonies we sampled for microscopic investigation varied from 2.5 to 17.5 m. A rough estimation suggests that their age ranges from 2.5 to 35 years. While the comparison between the age groups did not reveal difference in fungal composition, it is possible that younger sporophytes might exhibit different fungi composition in their fine roots. Additionally, seasonal changes could also change the species composition. For instance, the colonization of Mucoromycota endophytes in *Lycopodiella inundata* roots showed strong seasonal variation, with 86% colonization in autumn and only 14% in spring (Kowal et al. 2020). Our data is limited to a single collection time in August. Studying younger sporophytes and conducting collections at multiple time points would offer better insights into the relationship between *D. complanatum* sporophytes and their fungal partners.

#### 4.2.2. Pinus sylvestris

Considering a high host specificity of ECM symbiosis, especially increased by the high soil acidity and monodominance of the studied forest (Smith and Read 2008; Tederesoo et al. 2024), we expected *P. sylvestris* to have highest abundance of ECM fungi together with the lowest overall OTU diversity numbers. Compared to *D. complanatum*, *P. sylvestris* samples had significantly higher relative abundances of both symbiotrophic and ECM fungi along with the weaker association with saprotrophic fungi (Figs. 6a, 6b). The same pattern was observed in comparison with *V. myrtillus*, although it was not supported statistically.

The comparison of relative abundances per taxa (Figs. 4c, 4d, 4e) showed a significantly higher association of *P. sylvestris* only with genera *Russula* and *Tomentellopsis* compared to *V. myrtillus* and *D. complanatum*, respectively. Additionally, the analysis did not reveal differences in abundance of any ECM fungi between *P. sylvestris* and soil samples, most probably due to the high content of mycelia and spores of ECM fungi in the soil. Only the saprotrophic genus *Luellia* was more abundant in *P. sylvestris* fine roots than in the soil, and both *Luellia* and *Exophiala* were more abundant in *D. complanatum* samples, possibly playing an important role in root turnover of these species. Fortunately, no ASVs in the dataset were assigned to the genus *Heterobasidion* or its saprotrophic family Bondarzewiaceae, the widely distributed and most destructive disease agent of conifer trees, including *P. sylvestris* (Garbelotto and Gonthier 2013).

As expected, the calculated Hill numbers revealed that *P. sylvestris* had the lowest values for OTU richness and the diversity of common and dominant OTUs compared to the other sample types, even when considering only ECM fungi (Figs. 5a-c, Suppl. Figs. 8a-c).

#### 4.2.3. Vaccinium myrtillus

Despite the concept that ERM has the highest host-specificity among all types of mycorrhizae, *V. myrtillus* samples showed significantly higher values of OTU diversity than *P. sylvestris* samples (Fig. 7). Only for OTU richness calculated using only ECM fungi the confidence intervals overlapped with those for *P. sylvestris*, although they remained higher (Suppl. Fig. 7). Similar to *D. complanatum* and soil samples, this higher diversity of ECM fungi may be attributed to the presence of ECM fungi that also function as endophytes or sporophytes.

In our study, all fragments of *V. myrtillus* fine roots investigated microscopically showed the presence of dense hyphal coils inside the rhizodermal root cells, and these structures are characteristic for ERM (Figs. 7d, 7e). These coils could be formed by an association with Serendipitaceae (Sebacinales). Sebacinales were previously isolated from hyphal complexes of *V. myrtillus* collected in Europe and the ability of the genus *Serendipita* to colonise the roots of this species was even proved experimentally (Selosse et al. 2007; Vohník et al. 2016). Additionally, we found this family to be significantly more abundant in *V. myrtillus* compared to *P. sylvestris* root samples (Fig. 4d,). Interestingly, the order *Trechisporales* was more frequently found in *V. myrtillus* than in the other root samples, with its genus *Trechispora* being more abundant than in *D. complanatum*. This genus has been previously observed to form hyphal coils in the ericoid plants (Vohník et al. 2012). However, additionally to the intercellular structure, this fungus produces a hyphal sheath around the hair roots, which was not observed in our samples. The other genus potentially forming ERM, *Cladophialophora* (Herpotrichiellaceae, Chaetothyriales) (Allen et al. 2003), was also identified in our dataset, but did not show predominance in *V. myrtillus* samples. No other OTUs were assigned to the confirmed or putative ERM fungi listed in Leopold et al. (2016). Fungi that form ERM are believed to exclusively associate with *Ericaceae* and no other non-Ericaceae plants (Smith and Read 2008). This could explain why two out of three putative ERM families were predominantly found in *V. myrtillus* samples.

## 5. Conclusion

Both metabarcoding and microscopic approaches revealed differences in root-associated fungi between three closely growing, but phylogenetically distant plant species: the higher abundance of ECM was characteristic for *P. sylvestris* fine roots, while ERM was found predominant in *V. myrtillus* samples and neither of them, including AM, were dominant in *D. complanatum* samples. Metabarcoding analysis revealed that 66% of all identified taxa were found in all sample types. However, it also showed a significantly lower species diversity for *P. sylvestris* samples with the more abundant ECM fungi compared to other sample types. The ECM fungi associated with *P. sylvestris* fine roots were dominantly represented by *Russula* and *Tomentellopsis*. These results support the previous reports of high host specificity of ECM symbiosis. The fungal community associated with V. *myrtillus* root hairs was most similar to the one from *P. sylvestris*. However, *V. myrtillus* had a more diverse fungal community and two prevalent putative ERM fungal (classes Sebacinales and Trechisporales). No signs of AM were found by either method in any of the sample types. The abundant presence of Mucoromycota taxa, along with microscopy observations, suggests that Mucoromycota endophytes can inhabit the roots of *D. complanatum* sporophytes. However, the nature of this relationship requires further investigation. The absence of any type of mycorrhiza in *D. complanatum* sporophytes, substantially stronger development of root hairs and similarity to the soil samples together are pointing towards a low dependency from any known mycorrhiza type and an opportunistic kind of fungi-plant relationship. This is the first DNA sequencing study of fungi associated with *D. complanatum*. Additionally, by minimizing spatial variation through fine-scale sampling, here we contribute to the understanding of fungal interactions with plants from distant phylogenetic lineages in natural environments.

## Acknowledgements

We extend our sincere gratitude to the administration of Dzūkija National Park for their hospitality and support in data collection. Our appreciation also goes to Dr. Elke Seeber and our students for locating plots and recording vegetation data. *Vielen Dank* to Jan Woyzichovski for his expertise and invaluable assistance in preparing the light microscopy images. We are grateful to Oleg Shchepin for his contributions to data analysis and to Ulrich Möbius for his help with soil pH measurements. Special thanks go to Dr. Lorrie Maccario for preparing the DNA library and to Dr. Flavius Popa for aiding in the literature search. Finally, we are thankful to Greta Valvonyte for providing the photograph of *Diphasiastrum complanatum* roots (Suppl. Fig. 1a).

## Data availability statement

The sequencing data are available on NCBI (PRJNA1185013), the R and bash scripts of the data analysis, as well as created datasets, are available on GitHub: https://github.com/kuprinak/lithuania-metabar

## CRediT authorship contribution statement

Conceptualization: Martin Schnittler, Kristina Kuprina, Maria Sanchez Luque; Methodology: Kristina Kuprina, Maria Sanchez Luque, Heike Heklau; Formal analysis and investigation: Kristina Kuprina, Maria Sanchez Luque, Moana Wirth, Heike Heklau; Writing - original draft preparation: Kristina Kuprina; Writing - review and editing: Kristina Kuprina, Moana Wirth, Radvilė Rimgailė- Voicik, Manuela Bog, Martin Schnittler; Funding acquisition: Martin Schnittler; Supervision: Manuela Bog, Martin Schnittler.

## Abbreviations

RAF: root associated fungi
AM: arbuscular mycorrhiza
OTU: operational taxonomic unit
ECM: ectomycorrhiza
ERM: ericoid mycorrhiza
NTC: non-template control

## References

Abarenkov K, Nilsson RH, Larsson K-H, Taylor AFS, May TW et al (2023) The UNITE database for molecular identification and taxonomic communication of fungi and other eukaryotes: sequences, taxa and classifications reconsidered. Nucleic Acids Res 52(D1):D791–797. 10.1093/nar/gkad1039

Akhmetzhanova AA, Soudzilovskaia NA, Onipchenko VG, Cornwell WK et al. (2012) A rediscovered treasure: mycorrhizal intensity database for 3000 vascular plant species across the former Soviet Union. Ecol Arch E093–059. 10.1890/11-1749.1

Allen TR, Millar T, Berch SM, Berbee ML (2003) Culturing and direct DNA extraction find different fungi from the same ericoid mycorrhizal roots. New Phytol 160(1):255–272. 10.1046/j.1469-8137.2003.00885.x

Alzarhani AK, Clark DR, Underwood GJC, Ford H, Cotton TEA, Dumbrell AJ (2019) Are drivers of root-associated fungal community structure context specific? ISME J 13(5):1330–1344. 10.1038/s41396-019-0350-y

Andrews S (2010) FastQC: A Quality Control Tool for High Throughput Sequence Data [Online]. Available online at: https://www.bioinformatics.babraham.ac.uk/projects/fastqc/

Augustaitis A, Bytnerowicz A (2008) Contribution of ambient ozone to Scots pine defoliation and reduced growth in the Central European forests: A Lithuanian case study. Environ Pollut 155(3): 436–445. 10.1016/j.envpol.2008.01.042

Bardou P, Mariette J, Escudié F, Djemiel C, KloppC (2014) jvenn: an interactive Venn diagram viewer. BMC Bioinformatics 15:293. 10.1186/1471-2105-15-293

Beimforde C, Schäfer N, Dörfelt H, Nascimbene PC, Singh H, Heinrichs J, Reitner J, Rana RS, Schmidt AR (2011) Ectomycorrhizas from a Lower Eocene angiosperm forest. New Phytol 192(4):988–996. 10.1111/j.1469-8137.2011.03868.x

Benjamini Y, Hochberg Y (1995) Controlling the False Discovery Rate: A Practical and Powerful Approach to Multiple Testing. J R Stat Soc Series B Stat Method, 57(1):289–300. 10.1111/j.2517-6161.1995.tb02031.x

Bödeker ITM, Clemmensen KE, de Boer W, Martin F, Olson Å, Lindahl BD (2014) Ectomycorrhizal *Cortinarius* species participate in enzymatic oxidation of humus in northern forest ecosystems. New Phytologist 203: 245–256. 10.1111/nph.12791

Bolger AM, Lohse M, Usadel B (2014) Trimmomatic: A flexible trimmer for Illumina Sequence Data. Bioinformatics, btu170. 10.1093/bioinformatics/btu170

Bruchmann H (1898) Über die Prothallien und die Keimpflanzen mehrerer europäischer Lycopodien, und zwar über die von Lycopodium clavatum, L. annotinum, L. complanatum und L. selago. Perthes, Gotha, Germany. [In German] 10.5962/bhl.title.116256

Brundrett MC (2002) Coevolution of roots and mycorrhizas of land plants. New Phytol 154(2):275–304. 10.1046/j.1469-8137.2002.00397.x

Brundrett MC, Tedersoo L (2018) Evolutionary history of mycorrhizal symbioses and global host plant diversity. New Phytologist 220(4): 1108–1115. 10.1111/nph.14976

Bruns TD, Bidartondo MI, Taylor DL (2002) Host specificity in ectomycorrhizal communities: what do the exceptions tell us? Integr Comp Biol 42(2):352–9. 10.1093/icb/42.2.352

Bzdyk R, Sikora K, Studnicki M, Aleksandrowicz-Trzcińska M (2022) Communities of mycorrhizal fungi among seedlings of Scots pine (*Pinus sylvestris* L.) growing on a clearcut in microsites generated by different site-preparation methods. Forests 13(2): 353 10.3390/f13020353

Clay K, Schardl C (2002) Evolutionary origins and ecological consequences of endophyte symbiosis with grasses. Am Nat 160(4):99–127. 10.1086/342161

Clemmensen KE, Bahr A, Ovaskainen O, Dahlberg A, Ekblad A, Wallander H, Stenlid J, Finlay RD, Wardle DA, Lindahl BD (2013) Roots and associated fungi drive long-term carbon sequestration in boreal forest. Science 339(6127):1615–8. 10.1126/science.1231923

Dahlberg A, Finlay RD (1999) Suillus. In: Cairney JWG, Chambers SM (eds) Ectomycorrhizal Fungi: Key Genera in Profile. Springer Verlag, Berlin Heidelberg, pp. 33–64.

Davison J, Öpik M, Daniell TJ, Moora M, Zobel M (2011) Arbuscular mycorrhizal fungal communities in plant roots are not random assemblages. FEMS Microb Ecol 78(1):103–115. 10.1111/j.1574-6941.2011.01103.x

Delavaux C, Smith-Ramesh L, Kuebbing S (2017) Beyond nutrients: a meta-analysis of the diverse effects of arbuscular mycorrhizal fungi on plants and soils. Ecology 98(8):2111–2119. 10.1002/ecy.1892

Dickie IA, Koele N, Blum JD, Gleason JD, McGlone MS (2014): Mycorrhizas in changing ecosystems. Botany 92: 149–160. 10.1139/cjb-2013-0091

Edgar RC, Haas BJ, Clemente JC, Quince C, Knight (2011) UCHIME improves sensitivity and speed of chimera detection. Bioinformatics 27(16):2194–200. https://doi:10.1093/bioinformatics/btr38

Ewels P, Magnusson M, Lundin S, Käller M (2016) MultiQC: summarize analysis results for multiple tools and samples in a single report. Bioinformatics 32(19):3047–3048. 10.1093/bioinformatics/btw354

Frank B, Trappe JM (2005) On the nutritional dependence of certain trees on root symbiosis with belowground fungi (an English translation of A.B. Frank’s classic paper of 1885). Mycorrhiza 15:267–275. 10.1007/s00572-004-0329-y

Freeberg JA (1962) *Lycopodium prothalli* and their endophytic fungi as studied in vitro. Am J Bot 49(5):530–5. 10.2307/2439425

Garbelotto M, Gonthier P (2013) Biology, epidemiology, and control of *Heterobasidion* species worldwide. Annul Rev Phytopathol 51(1):39–59. 10.1146/annurev-phyto-082712-102225

Grime J, Mackey J, Hillier S et al (1987) Floristic diversity in a model system using experimental microcosms. Nature 328:420–422. 10.1038/328420a0

Gulbinas Z, Samuila M (2002) Integrated monitoring in small wooded catchments in Lithuania. Geol Q 46(1):81–97. https://gq.pgi.gov.pl/article/view/7957

Haynes W (2013) Wilcoxon Rank Sum Test. In: Dubitzky W, Wolkenhauer O, Cho KH, Yokota H (eds) Encyclopedia of Systems Biology. Springer, New York, NY. 10.1007/978-1-4419-9863-7_1185

Horn K, Franke T, Unterseher M, Schnittler M, Beenken L (2013) Morphological and molecular analyses of fungal endophytes of achlorophyllous gametophytes of *Diphasiastrum alpinum* (Lycopodiaceae). Am J Bot 100:2158–2174. 10.3732/ajb.1300011

Hsieh TC, Ma KH, Chao A (2022) iNEXT: Interpolation and Extrapolation for Species Diversity. R package version 3.0.0. http://chao.stat.nthu.edu.tw/wordpress/software_download/

Hutchison LJ (1999) *Lactarius*. In: Cairney JWG, Chambers SM (eds) Ectomycorrhizal Fungi: Key Genera in Profile. Springer Verlag, Berlin Heidelberg, pp 269–285

Ihrmark K, Bödeker IT, Cruz-Martinez K, Friberg H, Kubartova A, Schenck J, Strid Y, Stenlid J, Brandström-Durling M, Clemmensen KE, Lindahl BD (2012) New primers to amplify the fungal ITS2 region--evaluation by 454-sequencing of artificial and natural communities. FEMS Microbiol Ecol 82(3):666–77. 10.1111/j.1574-6941.2012.01437.x

James TY, Seifert KA (2017) Description of *Bifiguratus adelaidae*: The hunt ends for one of the “Top 50 Most Wanted Fungi.” Mycologia 109(3):361–362. 10.1080/00275514.2017.1372667

Karst J, Jones MD, Hoeksema JD (2023). Positive citation bias and overinterpreted results lead to misinformation on common mycorrhizal networks in forests. Nat Ecol Evol 7:501–511. 10.1038/s41559-023-01986-1

Kassambara A (2023) rstatix: Pipe-Friendly Framework for Basic Statistical Tests. R package version 0.7.2. https://CRAN.R-project.org/package=rstatix

Kemp E, Adam P, Ashford A (2003) Seasonal changes in hair roots and mycorrhizal colonization in *Woollsia pungens* (Cav.) F. Muell. (Epacridaceae). Plant and Soil 250:241–248. 10.1023/A:1022896014205.

Kennedy P, Izzo A, Bruns T (2003) There is high potential for the formation of common mycorrhizal networks between understorey and canopy trees in a mixed evergreen forest. J Ecol 91. 10.1046/j.1365-2745.2003.00829.x

Kiers ET, Duhamel M, Beesetty Y, Mensah JA, Franken O, Verbruggen E, Fellbaum CR, Kowalchuk GA, Hart MM, Bago A, Palmer TM, West SA, Vandenkoornhuyse P, Jansa J, Bücking H. (2011) Reciprocal rewards stabilize cooperation in the mycorrhizal symbiosis. Science 333(6044):880–2. 10.1126/science.1208473

Kjøller R, Olsrud M, Michelsen A (2010) Co-existing ericaceous plant species in a subarctic mire community share fungal root endophytes. Fungal Ecol 3: 205–214. 10.1016/j.funeco.2009.10.005

Kohout P (2017) Biogeography of Ericoid Mycorrhiza. In: Tedersoo L (ed) Biogeography of Mycorrhizal Symbiosis. Ecological Studies, vol 230. Springer, Cham. 10.1007/978-3-319-56363-3_9

Koide RT, Sharda JN, Herr JR, Malcolm GM (2008) Ectomycorrhizal fungi and the biotrophy-saprotrophy continuum. New Phytol 178(2):230–233. 10.1111/j.1469-8137.2008.02401.x

Kowal J, Arrigoni E, Serra J et al. (2020) Prevalence and phenology of fine root endophyte colonization across populations of *Lycopodiella inundata*. Mycorrhiza 30: 577–587. 10.1007/s00572-020-00979-3

Krings M, Taylor TN, Hass H, Kerp H, Dotzler N, Hermsen EJ (2007) Fungal endophytes in a 400-million-yr-old land plant: infection pathways, spatial distribution, and host responses. New Phytologist 174: 648–657. 10.1111/j.1469-8137.2007.02008.x

Kyaschenko J, Ovaskainen O, Ekblad A, Hagenbo A, Karltun E, Clemmensen KE, Lindahl BD (2019) Soil fertility in boreal forest relates to root-driven nitrogen retention and carbon sequestration in the mor layer. New Phytol 221(3):1492–1502. 10.1111/nph.15454

Lambers H, Shane MW, Cramer MD, Pearse SJ, Veneklaas EJ (2006) Root Structure and Functioning for Efficient Acquisition of Phosphorus: Matching Morphological and Physiological Traits. Ann Bot 98(4):693–713. 10.1093/aob/mcl114

Leopold D (2016) Ericoid fungal diversity: Challenges and opportunities for mycorrhizal research. Fungal Ecol 24:114–123. 10.1016/J.FUNECO.2016.07.004

Liu C, Cui Y, Li X, Yao M (2021) microeco: an R package for data mining in microbial community ecology. FEMS Microbiology Ecology 97(2): fiaa255. 10.1093/femsec/fiaa255

Love MI, Huber W, Anders S (2014). Moderated estimation of fold change and dispersion for RNA-seq data with DESeq2. Genome Biol 15:550. 10.1186/s13059-014-0550-8

Luo Y-H, Ma L-L, Seibold S, Cadotte MW, Burgess KS et al (2023) The diversity of mycorrhiza-associated fungi and trees shapes subtropical mountain forest ecosystem functioning. J Biogeogr 50: 715–729. 10.1111/jbi.14563

Majdi H, Damm E, Nylund J (2001) Longevity of mycorrhizal roots depends on branching order and nutrient availability. New Phytologist 150:195–202. 10.1046/J.1469-8137.2001.00065.X

Mandyam K, Jumpponen A (2008) Seasonal and temporal dynamics of arbuscular mycorrhizal and dark septate endophytic fungi in a tallgrass prairie ecosystem are minimally affected by nitrogen enrichment. Mycorrhiza 18:145–155. 10.1007/s00572-008-0165-6

McMurdie PJ, Holmes S (2014) Shiny-phyloseq: Web Application for Interactive Microbiome Analysis with Provenance Tracking. Bioinformatics (Oxford, England) 31(2):282–283. 10.1093/bioinformatics/btu616

Meddad-Hamza A, Hamza N, Neffar S, Beddiar A, Gianinazzi S, Chenchouni H (2017) Spatiotemporal variation of arbuscular mycorrhizal fungal colonization in olive (*Olea europaea* L.) roots across a broad mesic-xeric climatic gradient in North Africa. Sci Tot Environ 583:176–189. 10.1016/j.scitotenv.2017.01.049

National Land Service under the Ministry of Agriculture of the Republic of Lithuania (2016) Lithuanian National Atlas. Ministry of Agriculture of the Republic of Lithuania: Vilnius, Lithuania, Volume 1.

Nguyen NH, Song Z, Bates ST, Branco S, Tedersoo L, Menke J, Schilling JS, Kennedy PG (2016) FUNGuild: An open annotation tool for parsing fungal community datasets by ecological guild. Fungal Ecol 20:241–248. 10.1016/j.funeco.2015.06.006

Oinonen E (1967) Keltalieon (*Lycopodium complanatum* L.) itiöllinen uudistuminen etelä-suomessa kloonien laajuutta ja ikää koskevan tutkimuksen valossa (Sporal regeneration of ground pine (*Lycopodium complanatum* L.) in southern Finland in the light of the dimensions and the age of its clones). Acta For Fenn 83(3):1–85. [In Finnish] 10.14214/aff.7181

Olchowik J, Hilszczańska D, Studnicki M, Malewski T, Kariman K, Borowski Z (2021) Post-fire dynamics of ectomycorrhizal fungal communities in a Scots pine (*Pinus sylvestris* L.) forest of Poland. PeerJ 9:e12076. 10.7717/peerj.12076

Onguene NA, Kuyper TW (2001) Mycorrhizal associations in the rain forest of South Cameroon, For Ecol Manag 140(2–3):277–287. 10.1016/S0378-1127(00)00322-4

Op De Beeck M, Lievens B, Busschaert P, Declerck S, Vangronsveld J et al (2014) Comparison and Validation of Some ITS Primer Pairs Useful for Fungal Metabarcoding Studies. PLoS ONE 9(6):e97629. 10.1371/journal.pone.0097629

Öpik M, Moora M, Liira J, Kõljalg U, Zobel M, Sen R (2003) Divergent arbuscular mycorrhizal fungal communities colonize roots of *Pulsatilla spp*. in boreal Scots pine forest and grassland soils. New Phytol 160(3):581–593. 10.1046/j.1469-8137.2003.00917.x

Öpik M, Metsis M, Daniell TJ, Zobel M, Moora M (2009) Large-scale parallel 454 sequencing reveals host ecological group specificity of arbuscular mycorrhizal fungi in a boreonemoral forest. New Phytol 184(2):424–437. 10.1111/j.1469-8137.2009.02920.x

Orchard S, Standish RJ, Dickie IA, Renton M, Walker C, Moot D, Ryan MH (2017) Fine root endophytes under scrutiny: a review of the literature on arbuscule-producing fungi recently suggested to belong to the Mucoromycotina. Mycorrhiza 27(7):619–638. 10.1007/s00572-017-0782-z

Park SH, Eom AH (2007) Effects of mycorrhizal and endophytic fungi on plant community: a microcosm study. Mycobiology 35(4):186–90. 10.4489/MYCO.2007.35.4.186

Perez-Lamarque B, Petrolli R, Strullu-Derrien C, Strasberg D, Morlon H, Selosse MA, Martos F (2022) Structure and specialization of mycorrhizal networks in phylogenetically diverse tropical communities. Environ Microbiome 17(1):38. 10.1186/s40793-022-00434-0

Perez-Lamarque B, Laurent-Webb L, Bourceret A, Maillet L, Bik F, Cartier D, Labolle F, Holveck P, Epp D, Selosse MA (2023) Fungal microbiomes associated with Lycopodiaceae during ecological succession. Environ Microbiol Rep 15(2):109–118. 10.1111/1758-2229.13130

Perotto S, Girlanda M, Martino E (2002) Ericoid mycorrhizal fungi: some new perspectives on old acquaintances. Plant and Soil 244:41–53. 10.1023/A:1020289401610

Petrini O (1991) Fungal Endophytes of Tree Leaves. In: Andrews JH, Hirano SS (eds) Microbial Ecology of Leaves. Springer-Verlag, New York, pp 179–197. 10.1007/978-1-4612-3168-4_9

Posit Team (2024) RStudio: Integrated Development Environment for R. Posit Software, PBC, Boston, MA. http://www.posit.co

Quin D, Wang L, Han M, Wang J, Song H, Yan X, Duan X, Dong J (2018) Effect of an endophytic fungus *Umbelopsis dimorpha* on the secondary metabolites of host-plant *Kadsura agustifolia*. Front Microbiol 9:2845. 10.3389/fmicb.2018.02845

R Core Team (2024) R: A Language and Environment for Statistical Computing. R Foundation for Statistical Computing, Vienna, Austria. https://www.R-project.org.

Raven JA, Edwards D (2001) Roots: evolutionary origins and biogeochemical significance. J Exp Bot 52:381–401. 10.1093/jexbot/52.suppl_1.381

Redecker D, Kodner R, Graham LE (2000) Glomalean fungi from the Ordovician. Science 289: 1920–1921. 10.1126/science.289.5486.1920

Remy W, Taylor TN, Hass H, Kerp H (1994) Four hundred-million-year-old vesicular arbuscular mycorrhizae. Proc Natl Acad Sci USA. 91(25):11841–3. 10.1073/pnas.91.25.11841

Rho H, Hsieh M, Kandel S, Cantillo J, Doty S, Kim S (2017) Do Endophytes Promote Growth of Host Plants Under Stress? A Meta-Analysis on Plant Stress Mitigation by Endophytes. Microb Ecol 75.407–418. 10.1007/s00248-017-1054-3

Rice P, Longden I, Bleasby A (2000) EMBOSS: the European Molecular Biology Open Software Suite. Trends Genet 16(6):276–277. 10.1016/s0168-9525(00)02024-2

Rimington WR, Pressel S, Duckett JG, Field KJ, Read DJ, Bidartondo MI (2018) Ancient plants with ancient fungi: liverworts associate with early-diverging arbuscular mycorrhizal fungi. Proc Biol Sci 285(1888):20181600. 10.1098/rspb.2018.1600

Rimington WR, Duckett JG, Field KJ, Bidartondo MI, Pressel S (2020) The distribution and evolution of fungal symbioses in ancient lineages of land plants. Mycorrhiza 30:23–49. 10.1007/s00572-020-00938-y

Rodriguez RJ, Redman RS, Henson JM (2004) The Role of Fungal Symbioses in the Adaptation of Plants to High Stress Environments. Mitig Adapt Strateg Glob Chang 9:261–272. 10.1023/B:MITI.0000029922.31110.97

Rognes T, Flouri T, Nichols B, Quince C, Mahé F (2016) VSEARCH: a versatile open source tool for metagenomics. PeerJ 4:e2584. https://doi: 10.7717/peerj.2584

Rudawska M, Wilgan R, Janowski D, Iwański M, Leski (2018) Shifts in taxonomical and functional structure of ectomycorrhizal fungal community of Scots pine (*Pinus sylvestris* L.) underpinned by partner tree ageing. Pedobiologia 71: 20–30. 10.1016/j.pedobi.2018.08.003

Schmid E, Oberwinkler F (1993) Mycorrhiza-like interaction between the achlorophyllous gametophyte of *Lycopodium clavatum* L. and its fungal endophyte studied by light and electron microscopy. New Phytol 124: 69–81. 10.1111/j.1469-8137.1993.tb03798.x

Schroeder J, Martin J, Angulo D, Razo I, Barbosa J, Perea R, Sebastián-González E, Dirzo R (2019) Host plant phylogeny and abundance predict root-associated fungal community composition and diversity of mutualists and pathogens. J Ecol 107: 1557–1566. 10.1111/1365-2745.13166

Schwery O, Onstein RE, Bouchenak-Khelladi Y, Xing Y, Carter RJ, Linder HP (2915) As old as the mountains: the radiations of the Ericaceae. New Phytol 207(2):355–367. 10.1111/nph.13234

Selosse MA, Setaro S, Glatard F, Richard F, Urcelay C, Weiß M (2007) Sebacinales are common mycorrhizal associates of Ericaceae. New Phytol 174(4):864–878. 10.1111/j.1469-8137.2007.02064.x

Schnittler M, Horn K, Rico Kaufmann, Radvilė Rimgailė-Voicik, Anja Klahr, Manuela Bog, Jörg Fuchs, H. Wilfried Bennert (2019) Genetic diversity and hybrid formation in Central European club-mosses (*Diphasiastrum*, Lycopodiaceae) – New insights from cp microsatellites, two nuclear markers and AFLP. MBE 131:181–192. 10.1016/j.ympev.2018.11.001

Sietiö OM, Tuomivirta T, Santalahti M, Kiheri H, Timonen S, Sun H, Fritze H, Heinonsalo J (2018) Ericoid plant species and *Pinus sylvestris* shape fungal communities in their roots and surrounding soil. New Phytol 218(2):738–751. http://ddoi.org/10.1111/nph.15040

Simard SW, Beiler KJ, Bingham MA, Deslippe JR, Philip LJ, Teste FP (2012) Mycorrhizal networks: mechanisms, ecology and modelling. Fungal Biol Rev 26:39–60. 10.1016/j.fbr.2012.01.001

Slowikowski K (2024) ggrepel: Automatically Position Non-Overlapping Text Labels with ’ggplot2’. https://ggrepel.slowkow.com/. https://github.com/slowkow/ggrepel

Smith G, Finlay R, Stenlid J, Vasaitis R, Menkis A (2017) Growing evidence for facultative biotrophy in saprotrophic fungi: data from microcosm tests with 201 species of wood-decay basidiomycetes. The New Phytol 215(2):747–755. 10.1111/nph.14551

Smith SE, Read DJ (2008) Mycorrhizal symbiosis, 3rd edn. Amsterdam; Boston: Academic Press. 10.2136/sssaj2008.0015br

Spalding B, Duxbury JM, Stone EL (1975) *Lycopodium* fairy rings: effect on soil respiration and enzymatic activities. Soil Sci Soc Am J 39(1):65–70. 10.2136/sssaj1975.03615995003900010020x

Spessard EA (1917) Prothallia of *Lycopodium* in America. Bot Gaz 63:66–76. 10.1086/331967

State Forest Service under the Ministry of Environment (2023) Forest Cadastre Data [Valstybinė miškų tarnyba prie Aplinkos ministerijos. 2023. Miškų kadastro duomenys] [accessed 2023-05-05]. Source: https://www.geoportal.lt/map#portalAction=openService&serviceUrl=https%3A%2F%2Fwww.geoportal.lt%2Fmapproxy%2Fvmt_mkd%2FmapServer

Stone J, Polishook JD, and White JF (2004) Endophytic fungi. In: Mueller GM, Bills GF, White JF (eds) Biodiversity of Fungi, Amsterdam, Elsevier, pp 241–270. 10.1016/B978-012509551-8/50015-5

Strullu-Derrien C, Selosse MA, Kenrick P, Martin FM (2018) The origin and evolution of mycorrhizal symbioses: from palaeomycology to phylogenomics. New Phytol 220(4):1012–1030. 10.1111/nph.15076

Sun H, Terhonen E, Kovalchuk A, Tuovila H, Chen H, Oghenekaro AO,Heinonsalo J, Kohler A, Kasanen R, Vasander H, Asiegbu FO (2016) Dominant treespecies and soil type affect the fungal community structure in a boreal peatlandforest. Appl Environ Microbiol 82:2632–2643. 10.1128/AEM.03858-15

Taniguchi T, Kitajima K, Douhan GW et al (2018) A pulse of summer precipitation after the dry season triggers changes in ectomycorrhizal formation, diversity, and community composition in a Mediterranean forest in California, USA. Mycorrhiza 28:665–677. 10.1007/s00572-018-0859-3

Tedersoo L, Bahram M, Põlme S, Kõljalg U, Yorou NS, Wijesundera R, VillarrealRuiz L, Vasco-Palacios AM et al (2014) Global diversity and geography of soil fungi. Science 346(6213):1256688. 10.1126/science.1256688

Tedersoo L, Bahram M, Toots M, Diédhiou AG, Henkel TW, Kjøller R, Morris MH, Nara K, Nouhra E, Peay KG, Põlme S, Ryberg M, Smith ME, Kõljalg U (2012) Towards global patterns in the diversity and community structure of ectomycorrhizal fungi. Mol Ecol 21(17):4160–70. 10.1111/j.1365-294X.2012.05602.x

Tedersoo L, Brundrett M (2017) Evolution of ectomycorrhizal symbiosis in plants. Ecological Studies 230:407–467. 10.1007/978-3-319-56363-3_19

Tedersoo L, Drenkhan R, Abarenkov K, Anslan S, Bahram M, Bitenieks K, Buegger F, Gohar D, Hagh-Doust N, Klavina D, Makovskis K, Zusevica A, Pritsch K, Padari A, Põlme S, Rahimlou S, Rungis D, Mikryukov V (2024) The influence of tree genus, phylogeny, and richness on the specificity, rarity, and diversity of ectomycorrhizal fungi. Environ Microbiol Rep 16(2):e13253. 10.1111/1758-2229.13253

Tedersoo L, Jairus T, Horton BM, Abarenkov K, Suvi T, Saar I, Kõljalg U (2008) Strong host preference of ectomycorrhizal fungi in a Tasmanian wet sclerophyll forest as revealed by DNA barcoding and taxon-specific primers. New Phytol 180(2):479–490. 10.1111/j.1469-8137.2008.02561.x

Tedersoo L, Liiv I, Kivistik PA, Anslan S, Kõljalg U, Bahram M (2016) Genomics and metagenomics technologies to recover ribosomal DNA and single-copy genes from old fruit-body and ectomycorrhiza specimens. MycoKeys 13:1–20. 10.3897/mycokeys.13.8140

Tedersoo L, May TW, Smith ME (2010) Ectomycorrhizal lifestyle in fungi: global diversity, distribution, and evolution of phylogenetic lineages. Mycorrhiza 20(4):217–63. 10.1007/s00572-009-0274-x

Tedersoo L, Smith MS (2013) Lineages of ectomycorrhizal fungi revisited: Foraging strategies and novel lineages revealed by sequences from belowground. Fungal Biol Rev 27:83–99. 10.1016/j.fbr.2013.09.001

Teste F, Jones MD, Dickie IA (2020) Dual-mycorrhizal plants: their ecology and relevance. New Phytol 225(5):1835–1851. 10.1111/nph.16190

Toju H, Yamamoto S, Sato H, Tanabe AS (2013) Sharing of diverse mycorrhizal and root-endophytic fungi among plant species in an oak-dominated cool-temperate forest. PLoS One 8(10):e78248. 10.1371/journal.pone.0078248

Treu R, Laursen GA, Stephenson SL, Landolt JC, Densmore R (1996) Mycorrhizae from Denali National Park and Perserve, Alaska. Mycorrhiza 6: 21–29.

Tukey JW (1949) Comparing individual means in the analysis of variance. Biometrics 5(2):99–114. 10.2307/3001913

Vandenkoornhuyse P, Husband R, Daniell TJ, Watson IJ, Duck JM, Fitter AH, Young JP (2002) Arbuscular mycorrhizal community composition associated with two plant species in a grassland ecosystem. Mol Ecol 11(8):1555–64. 10.1046/j.1365-294x.2002.01538.x

Verbruggen E, van der Heijden MG, Weedon JT, Kowalchuk GA, Röling WFM (2012) Community assembly, species richness and nestedness of arbuscular mycorrhizal fungi in agricultural soils. Mol Ecol 21:2341–2353. 10.1111/j.1365-294X.2012.05534.x

Veresoglou SD, Caruso T, Rillig MC (2013) Modelling the environmental and soil factors that shape the niches of two common arbuscular mycorrhizal fungal families. Plant and Soil 368:507–518. 10.1007/s11104-012-1531-x

Vohník M, Sadowsky JJ, Kohout P, Lhotáková Z, Nestby R et al (2012) Novel Root-Fungus Symbiosis in Ericaceae: Sheathed Ericoid Mycorrhiza Formed by a Hitherto Undescribed Basidiomycete with Affinities to Trechisporales. PLOS ONE 7(6):e39524. 10.1371/journal.pone.0039524

Vohník M, Pánek M, Fehrer J et al. (2016) Experimental evidence of ericoid mycorrhizal potential within Serendipitaceae (Sebacinales). Mycorrhiza 26:831–846. 10.1007/s00572-016-0717-0

Vohník M (2020) Ericoid mycorrhizal symbiosis: theoretical background and methods for its comprehensive investigation. Mycorrhiza 30:671–695. 10.1007/s00572-020-00989-1

Walker JF, Aldrich-Wolfe L, Riffel A, Barbare H, Simpson NB, Trowbridge J, Jumpponen A (2011) Diverse Helotiales associated with the roots of three species of Arctic Ericaceae provide no evidence for host specificity. New Phytol 191(2):515–527. 10.1111/j.1469-8137.2011.03703.x

Wang B, Qiu YL (2006) Phylogenetic distribution and evolution of mycorrhizas in land plants. Mycorrhiza 16:299–363. 10.1007/s00572-005-0033-6

Waud M, Busschaert P, Ruyters S, Jacquemyn H, Lievens B (2014) Impact of primer choice on characterization of orchid mycorrhizal communities using 454 pyrosequencing. Mol Ecol Res 14(4): 679–699. 10.1111/1755-0998.12229

Weng JK, Li X, Stout J, Chapple C (2008) Independent origins of syringyl lignin in vascular plants. Proc Natl Acad Sci USA 105(22):7887–92. 10.1073/pnas.0801696105

Weng JK, Chapple C (2010) The origin and evolution of lignin biosynthesis. New Phytol 187:273–285. 10.1111/j.1469-8137.2010.03327.x

Whittier DP (1977) Gametophytes of Lycopodium obscurum as grown in axenic culture. Canad J Bot 55:563–567. 10.1139/b77-067

Wickham H (2016) ggplot2: Elegant Graphics for Data Analysis. Springer-Verlag New York. https://ggplot2.tidyverse.org

Wikström N, Kenrick P (2001) Evolution of Lycopodiaceae (Lycopsida): Estimating divergence time from rbcL gene sequences by use of nonparametric rate smoothing. Mol Phylogenet Evol 19:177–186. 10.1006/mpev.2001.0936

Winther JL, Friedman WE (2008) Arbuscular mycorrhizal associations in Lycopodiaceae. New Phytol 177(3):790–801. 10.1111/j.1469-8137.2007.02276.x

Zhang J, Quan C, Ma L et al (2021) Plant community and soil properties drive arbuscular mycorrhizal fungal diversity: A case study in tropical forests. Soil Ecol Lett 3:52–62. 10.1007/s42832-020-0049-z

