## Supplementary material for "Phylogenetically distant but cohabiting: Fungal communities of fine roots in *Diphasiastrum complanatum*, *Pinus sylvestris*, and *Vaccinium myrtillus* in a Lithuanian pine forest": Suppl.

Supplementary Table 1. Description of 19 plots included to the study.

| Plot name | Coordinates  (Latitude, Longitude) | Forest age, years | Forest height, m | *Diphasiastrum complanatum* colony diameter, m | Soil pH  (2021/2023) |
| --- | --- | --- | --- | --- | --- |
| 30A | 54.11406, 24.30972 | 101 | 27.12 | 15.0 | 3.39/3.97 |
| 2 | 54.12325, 24.28862 | 115 | 28.00 | 17.5 | 3.31/3.28 |
| 4 | 54.12601, 24.28183 | 115 | 27.00 | 8.5 | 3.32/3.34 |
| 5 | 54.12457, 24.27913 | 120 | 27.00 | 4.5 | 2.89/NA |
| 6 | 54.12301, 24.27845 | 105 | 28.00 | 5.5 | 3.57/NA |
| 9 | 54.12190, 24.27683 | 116 | 25.78 | 9.0 | 3.13/3.06 |
| 10 | 54.12355, 24.28391 | 115 | 28.00 | 16.5 | 3.16/NA |
| 78 | 54.10849, 24.30301 | 81 | 25.78 | 9.0 | 3.41/3.51 |
| 76 | 54.10770, 24.30784 | 91 | 24.31 | 11.0 | 3.42/3.61 |
| 22 | 54.10401, 24.31006 | 91 | 24.31 | 8.5 | 3.33/3.37 |
| 23 | 54.10384, 24.31421 | 121 | 24.69 | 25.0 | 3.39/NA |
| 24 | 54.09533, 24.32623 | 105 | 27.00 | 2.0 | 3.24/NA |
| 16 | 54.10704, 24.30510 | 76 | 20.70 | 5.0 m line | 3.77/NA |
| 17 | 54.10425, 24.31343 | 121 | 24.69 | 6.5 | 3.40/NA |
| 26 | 54.10326, 24.35016 | 96 | 25.19 | 10.0 | 3.27/NA |
| 27 | 54.09906, 24.35385 | 120 | 27.00 | 2.5 | 3.24/3.20 |
| 13 | 54.10168, 24.35578 | 101 | 28.14 | 17.0 | 3.24/3.35 |
| 30B | 54.10388, 24.35548 | 126 | 25.67 | 6.0 | 3.45/3.40 |
| 31 | 54.10306, 24.35635 | 116 | 25.78 | 5.0 | 3.33/NA |
| **Average** | **-** | **106** | **26.01** | **9.94** | **3.33/3.41** |

Supplementary Table 2. Average percentage (± sd) of vegetation coverage across five strata, along with the corresponding plant/lichen species found in the 19 investigated 5 x 5 m plots.

| Trees  (> 3 m tall)  31.2 ± 24 | Shrubs  (0.5-3 m)  11.5 ± 8.8 | Subshrubs and herbs  (< 0.5 m)  17.9 ± 12 | Mosses and lichens  75.4 ± 17 | Wood, stones, bare soil  14.9 ± 12.0 |
| --- | --- | --- | --- | --- |
| *Pinus sylvestris*  28.0 ± 25.0  *Betula pendula*  6.1 ± 9.7  *Quercus robur*  0.6 ± 1.5  *Picea abies*  0.3 ± 1.2  *Prunus serotina*  0.2 ± 0.4  *Sorbus aucuparia*  0.08 ± 0.3 | *Juniperus communis*  9.3 ± 1.0  *Frangula alnus*  0.6 ± 0.7 | *Vaccinium vitis-idaea*  6.6 ± 12  *Diphasiastrum complanatum*  6.3 ± 5.3  *Vaccinium myrtillus*  4.6 ± 5.7  *Melampyrum sylvaticum*  0.8 ± 1.5  *Avenella flexuosa*  0.7 ± 1.2  other species* (25)  0.1 ± 0.4 | *Hylocomium splendens*  35.6 ± 23.0  *Pleurozium schreberi*  28.4 ± 24.0  *Dicranum polysetum*  8.9 ± 11.0  *Polytrichum commune*  2.4 ± 3.1  *Cladonia rangiferina*  2.0 ± 4.0  other species** (11)  0.3 ± 4.2 | - |

* *Chimaphila umbellata*, *Solidago virgaurea, Carex ericetorum, Calamagrostis villosa, Hieracium umbellatum, Agrostis vinealis, Dryopteris filix-mas, Hieracium lachenalii, Lycopodium clavatum, Rumex acetosella, Goodyera repens, Rubus idaeus, Anthoxanthum odoratum, Carex leporina, Corynephorus canescens, Danthonia decumbens, Hieracium pilosella, Hypericum maculatum, Thymus serpyllum, Trientalis europaea, Veronica officinalis, Calamagrostis epigeios, Nardus stricta, Pyrola rotundifolia, Monotropa hypopitys*

*** Dicranum scoparium, Ptilium crista-castrensis, Polytrichum juniperinum, Aulacomnium palustre, Ptilidium ciliare, Sphagnum girgensohnii, Mnium hornum, Polytrichum formosum, Dicranum sp., Cetraria islandica*

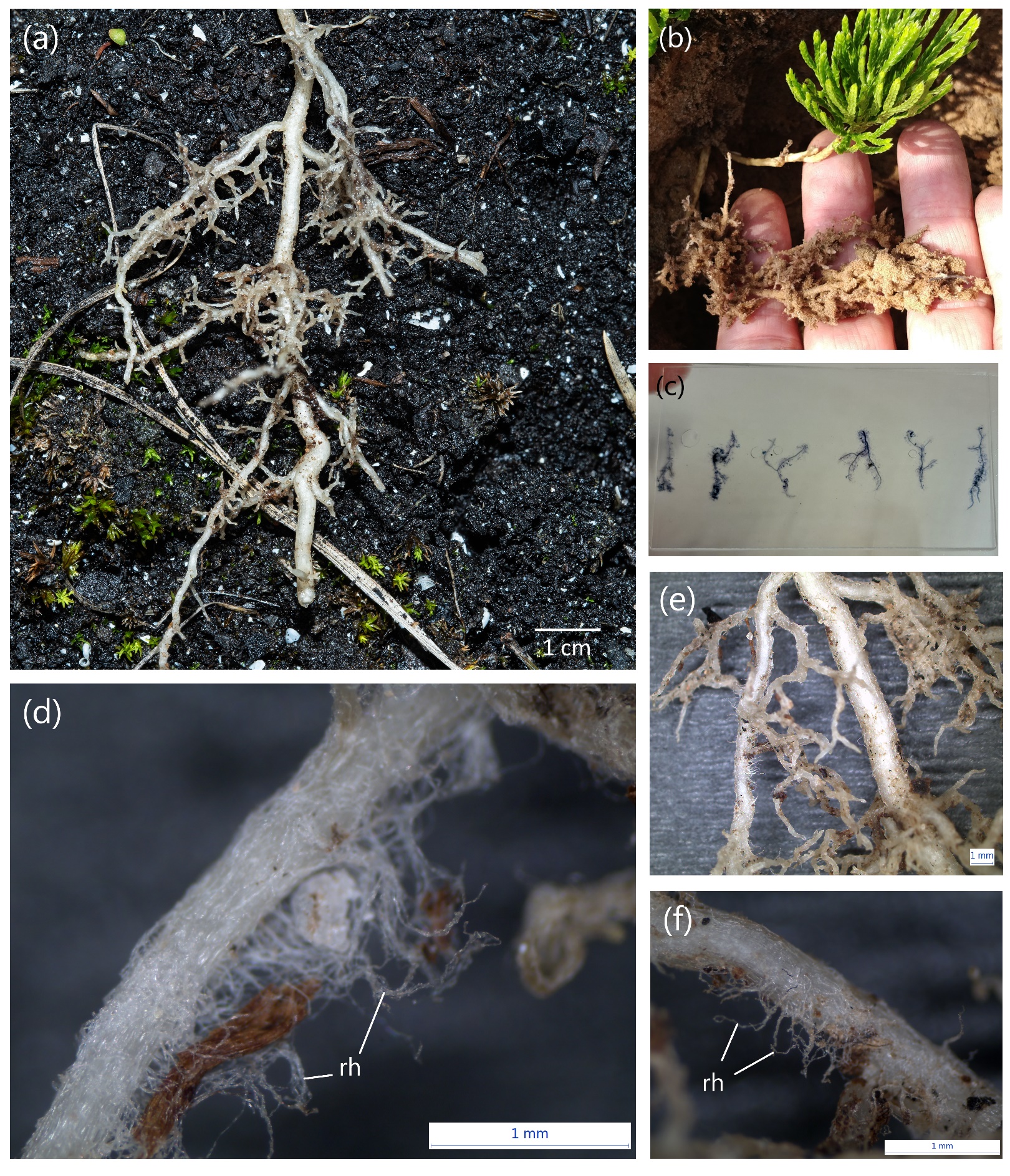

Supplementary Figure 1. (a-f) Roots of sporophytes from *Diphasiastrum complanatum* with a dense cover of root hairs [rh].

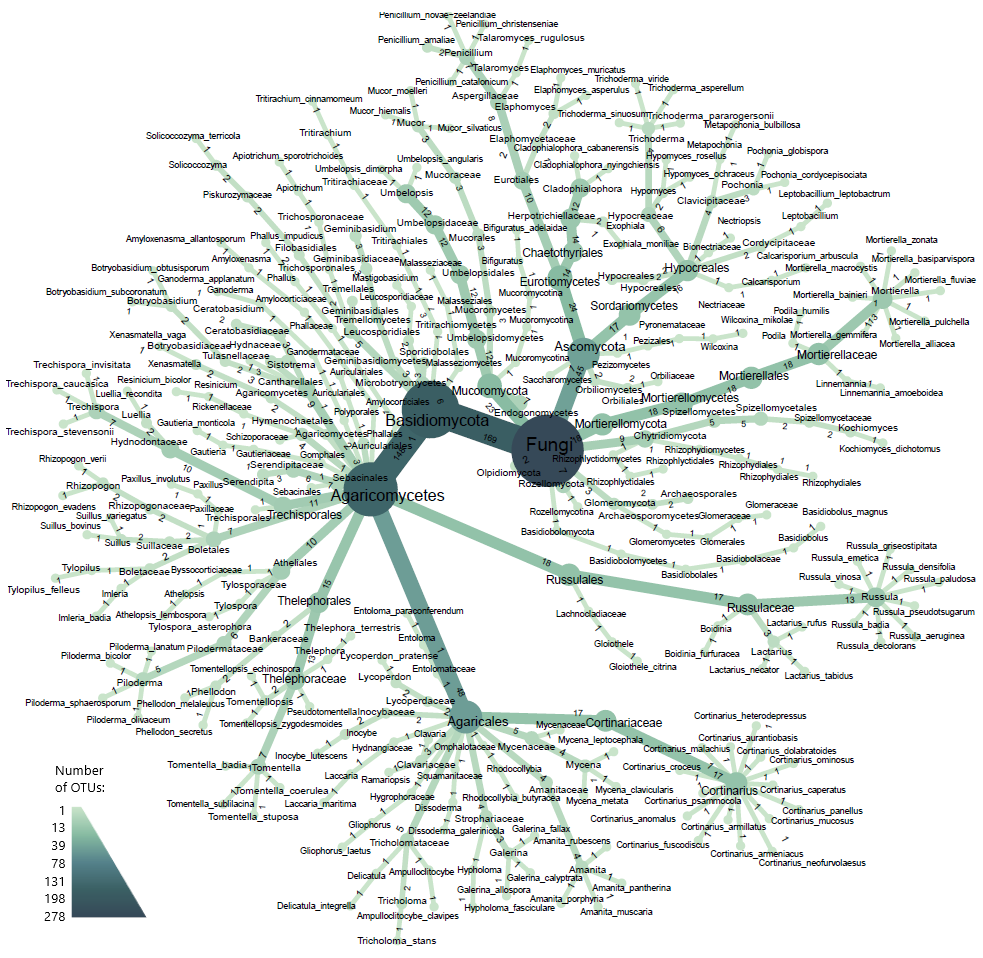

Supplementary Figure 2. A heat tree illustrating the diversity of all fungal taxa identified within 278 OTUs through metabarcoding analysis of 76 samples of soil or fine roots of *Pinus sylvestris*, *Vaccinium myrtillus*, and *Diphasiastrum complanatum* in 19 studied plots. The size and the colour of the nodes, as well as the numbers in the branches, indicate the number of corresponding OTUs.

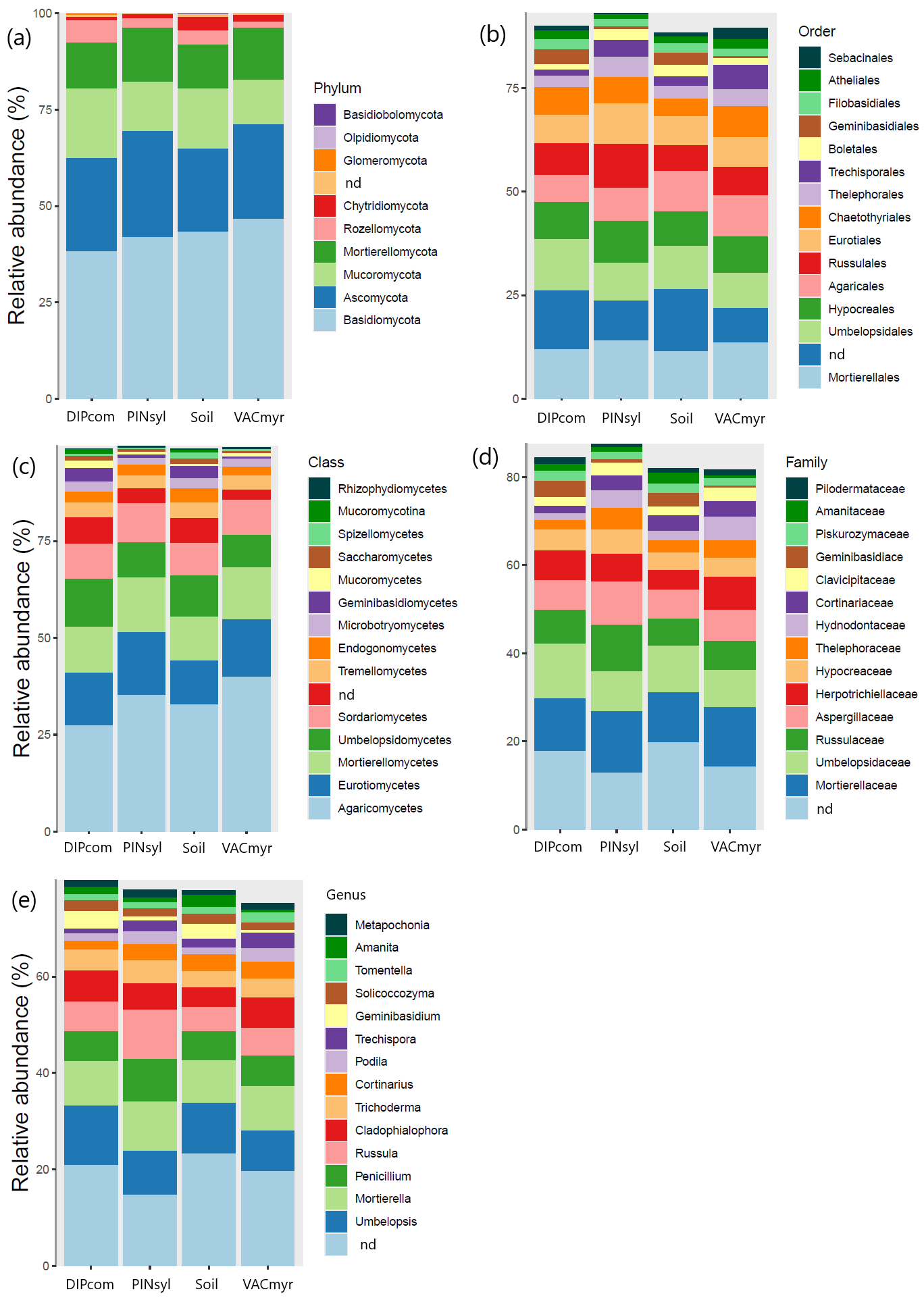

Supplementary Figure 3. Relative abundance of normalised reads of 278 OTUs assigned to nine phyla (a), 15 most abundant classes (b), orders (c), families (d) and genera (e) across samples of soil, fine roots of *Diphasiastrum complanatum* (DIPcom), *Pinus sylvestris* (PINsyl), and *Vaccinium myrtillus* (VACmyr). Samples were collected in 19 plots for each sample type (n = 74 in total).

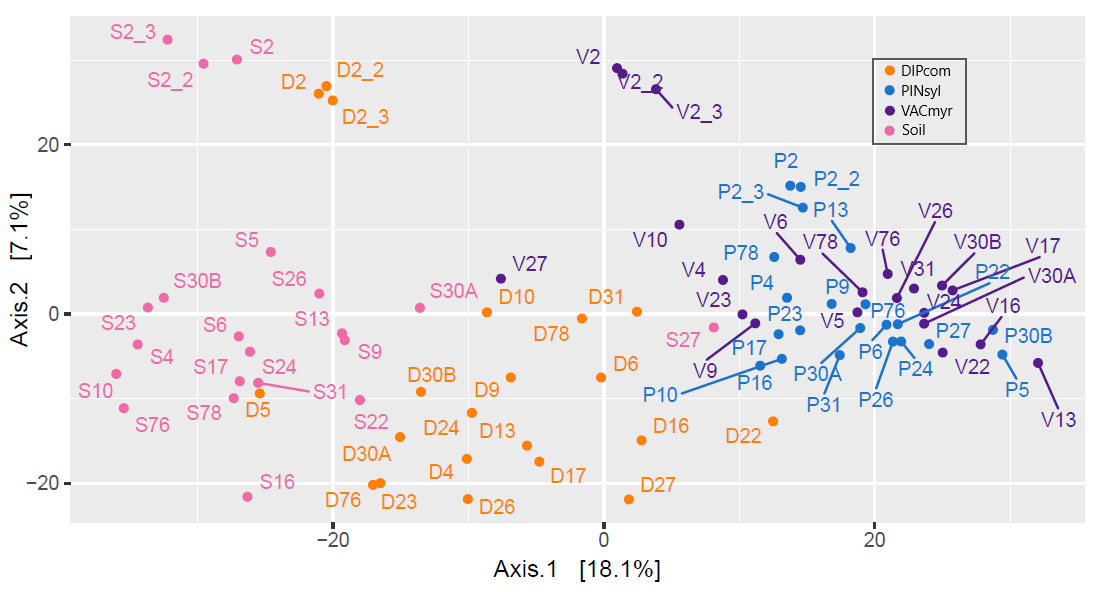
Supplementary Figure 4. Principal Coordinate Analysis (PCoA) ordination diagram generated from Euclidean distances between normalized counts of 278 OTUs found in soil, fine roots of *Diphasiastrum complanatum* sporophytes (DIPcom), *Pinus sylvestris* (PINsyl), and *Vaccinium myrtillus* (VACmyr). Samples were collected in 19 plots (83 samples in total); technical replicates (n = 8) contain underscores in the corresponding labels.

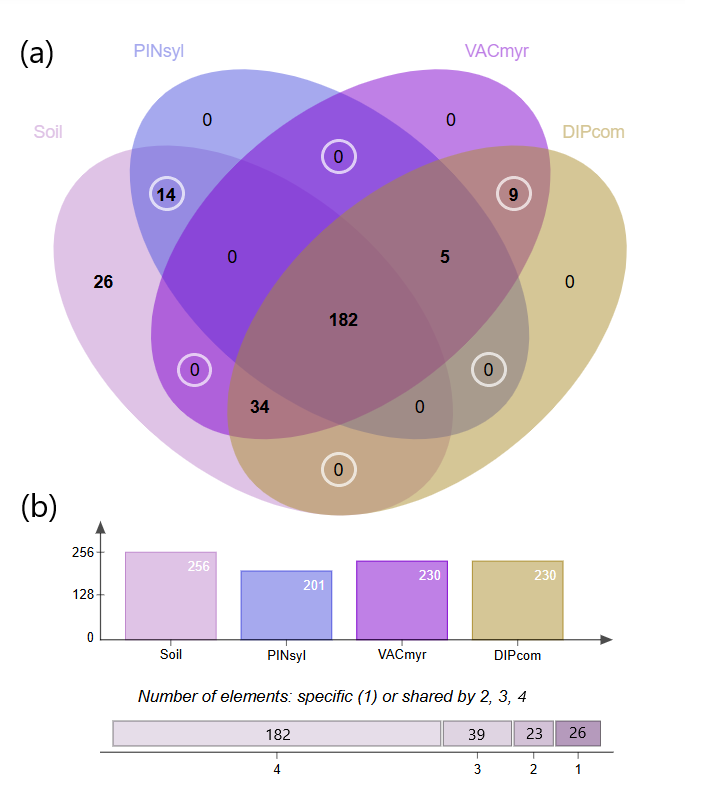

Supplementary Figure 5. (a) Venn diagram showing the number of unique and shared OTUs and (b) total numbers of OTUs across samples of soil or fine roots of *Diphasiastrum complanatum* sporophytes (DIPcom), *Vaccinium myrtillus* (VACmyr), and *Pinus sylvestris* (PINsyl) collected from 19 studied plots (76 samples and 278 ASVs, in total). White circles indicate pairwise comparisons between sample types.

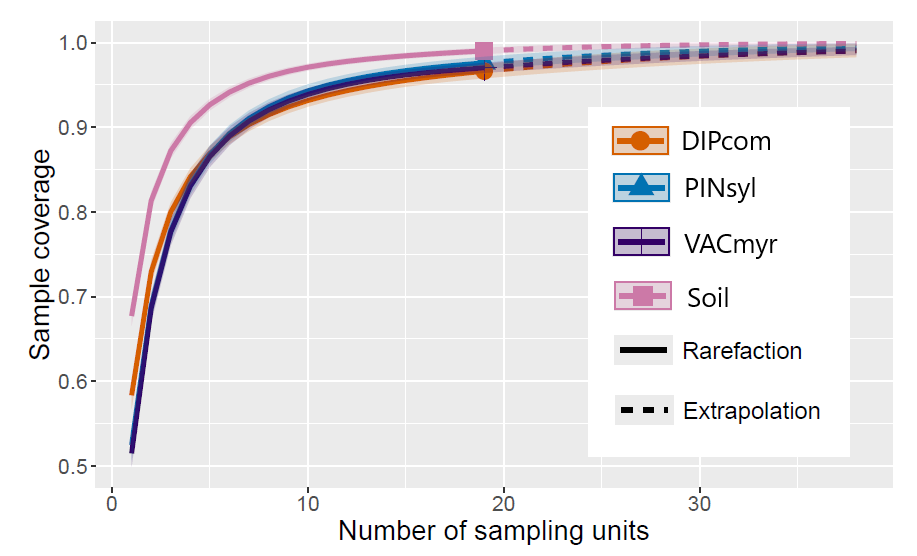

Supplementary Figure 6. Rarefaction and extrapolation curves with 95% confidence intervals for 278 OTUs detected in soil and fine roots of *Diphasiastrum complanatum* sporophytes (DIPcom) *Pinus sylvestris* (PINsyl), *Vaccinium myrtillus* (VACmyr), and collected in 19 plots (76 samples in total).

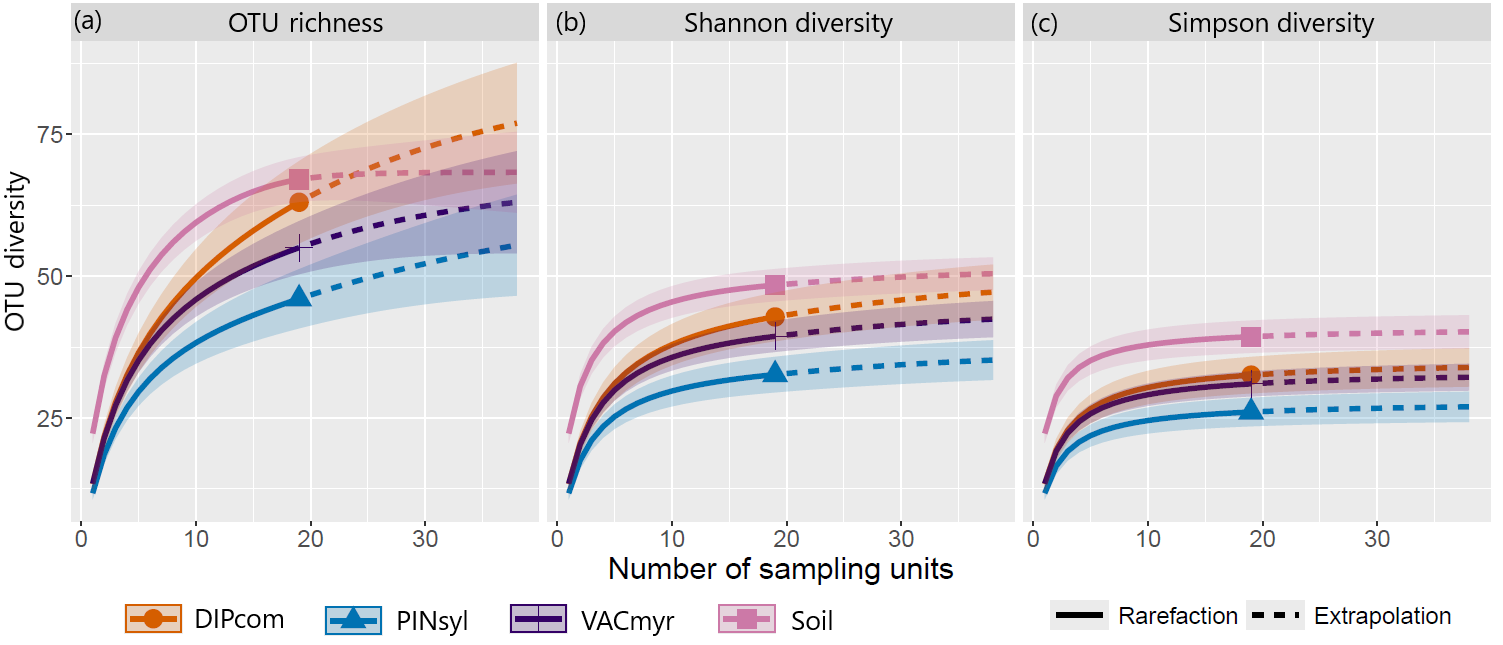

Supplementary Figure 7. Rarefaction and extrapolation curves with 95% confidence intervals of three Hill numbers describing OTU diversity: (a) OTU richness, (b) Shannon diversity, (c) Simpson diversity. The numbers are calculated using normalized counts of 86 OTUs assigned to ectomycorrhizal fungi for soil or fine roots of *Diphasiastrum complanatum* sporophytes (DIPcom) *Pinus sylvestris* (PINsyl), *Vaccinium myrtillus* (VACmyr), and collected in 19 plots (76 samples in total).
